# Coupling Mechanical Regulation with Biochemical Reaction-Diffusion Circuits Yields Robust Self-Organized Pattern Formation

**DOI:** 10.64898/2026.05.23.727407

**Authors:** Satoshi Toda, Guoye Guan, Mai Tambo, Hiromi Morita, Wendell A. Lim

## Abstract

Cell-cell signaling circuits that combine local self-activation with long-range inhibition have long been proposed as a theoretical mechanism sufficient to generate cellular patterns, such as spots and stripes. Here we construct synthetic pattern-forming circuits, implementing local self-activation (positive feedback) using juxtacrine synNotch receptor interactions, and implementing long-range inhibition using diffusible competitor molecules. While combining local positive feedback with long-range inhibition leads to more spatially heterogeneous cell states, these synthetic circuits do not robustly lead to well defined patterns. We find, however, that if we couple these reaction-diffusion circuits with induction of genes that regulate cell mechanics – such Cadherin molecules that promote local cell adhesion and sorting – we observed the emergence of much more well-defined cellular patterns. Theoretical analysis indicates while reaction-diffusion circuits can be sufficient to generate patterns under precisely balanced parameter conditions, the close coupling of cell mechanical/sorting significantly increases the robustness of pattern formation (the parameter space yielding patterns). Thus, circuits that closely integrate signaling and mechanical changes may underlie many evolved morphogenic pattern formation systems.

## Introduction

One of the hallmarks that distinguishes living organisms from non-living entities is self-organization. Cells in multicellular organisms interact with one another to robustly self-organize into diverse spatial patterns during development. Chemical reaction-diffusion systems have long been proposed as a theoretical framework to explain how interacting agents can generate spatial patterns from an initially homogeneous status (*1*). In particular, when slowly-diffusing activators promote their own production through short-range positive feedback, while simultaneously inducing rapidly-diffusing inhibitors to generate long-range negative feedback, the homogeneous state becomes unstable and endogenous fluctuations can drive self-organization into periodic patterns such as spots and stripes. The concept of reaction-diffusion systems has been extended into a more general framework of local autoactivation-lateral inhibition (LALI) to explain multicellular biological pattern formation (*2–4*) (**Fig. 1A**). Genetic analyses in model organisms support a central role of morphogen-based LALI-circuits in generating spatial tissue patterns such as hair follicles, digits, and fingerprints (*5–7*).

**Fig. 1.**
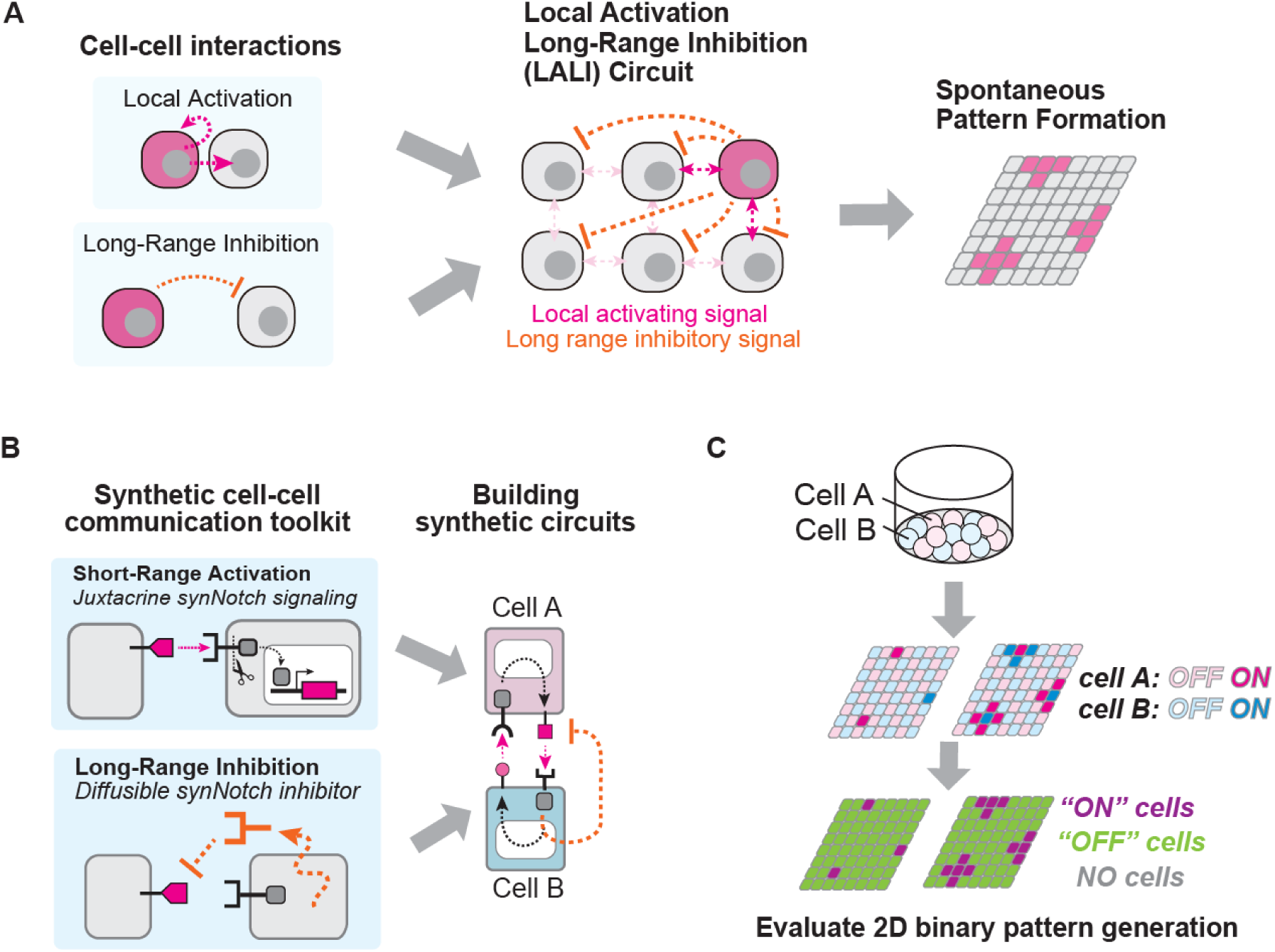
Synthetically exploring the design space of multi-cellular pattern forming reaction-diffusion circuits. (A) Schematics of spontaneous pattern formation by LALI (Local Activation and Long-Range Inhibition) circuit in multicellular systems. (B) Schematics of synNotch-based LALI circuit construction. synNotch ligand-receptor interactions mediate synthetic juxtacrine signaling that induces target gene expression, whereas secreted synNotch inhibitors function as diffusible inhibitory signals. Their combination enables construction of synthetic LALI circuits in mammalian cell culture. (C) Experimental workflow. 20000 cells of each Cell A and Cell B were cocultured in non-treat 96 well plates for 72 hr. Spatial patterns of induced fluorescent reporters were imaged by confocal microscope and processed to classify ON, OFF, and NO cell regions (magenta, green, and gray, respectively), to evaluate synthetic pattern generation.

There is growing interest in being able to engineer synthetic developmental systems, to both evaluate and refine our fundamental understanding of the design principles underlying development, and to move towards the predictive design of novel or modified tissues (*8–12*). Thus, a fundamental question is whether we can construct synthetic pattern forming systems using LALI based circuits. While LALI morphogen circuits clearly play a role in many native pattern forming systems, are they sufficient to yield patterns in a highly robust and reproducible manner? Many theoretical studies have characterized the potential of LALI circuits to drive pattern formation and the required system parameters (*13, 14*), but few experimental reconstitution studies have been done (*15, 16*). In mammalian systems, synthetic reaction diffusion circuits have been reconstructed using the native morphogens Nodal and Lefty (*17*). This system yielded some degree of pattern formation, but relied on native morphogens with evolved features (e.g. distinct diffusion ranges) that might be optimized for behaviors like pattern formation. Thus, a systematic analysis of circuit features that are necessary and sufficient to yield robust synthetic cell patterns remains largely unexplored.

Here we take advantage of recent advances in synthetic receptor technologies to construct completely synthetic LALI circuit in mammalian cells using non-natural “morphogens,” their receptors, and their inhibitors, providing an empirical framework to test what genetic circuits are sufficient to drive tissue patterning (*18, 19*). More specifically synthetic Notch (synNotch) receptors allow the construction of engineered juxtacrine cell-cell signaling through artificial ligand-receptor interactions, which can be linked to the induction of user-defined target genes (**Fig. 1B**) (*20*). SynNotch systems thus allow construction of a short-range positive feedback circuit by inducing the expression of synNotch ligand in response to synNotch activation. In addition, synNotch systems enable building a long-range negative feedback circuit by inducing expression of a competing high affinity nanobody that blocks the synNotch ligand, thus acting as a diffusible inhibitor (*18*). Combining these feedback circuits, we are able to build synthetic reaction diffusion circuits in mammalian cells with systematically altered circuit parameters (**Fig. 1B**).

By evaluating the 2D ON-OFF spatial patterns resulting from these circuits (**Fig. 1C**), we found that the synthetic LALI reaction-diffusion circuits alone were not sufficient to yield robust, well-defined multicellular patterns. Cell fate bifurcation and patterning was highly sensitive to the fine-tuned strength of negative feedback, indicating limited robustness. We found, however, that we could significantly improve both pattern definition and robustness, by coupling the LALI synthetic reaction-diffusion circuits with induction of cell adhesion. By integrating the morphogen signaling circuits with a mechanical change of increased cell adhesion, we observed well-defined multi-spot patterns emerging across a much broader range of parameter space. To comprehensively analyze the effects of induced cell adhesion, we developed an agent-based computational model, which revealed that coupling of cell adhesion to reaction-diffusion circuits is predicted to substantially expanded the parameter space supporting pattern formation in many different LALI circuit implementations. Together, these results demonstrate that integrating biochemical (morphogen) signaling with induced mechanical regulation enables the construction of far more robust genetic programs for predictable and tunable multicellular pattern formation.

### Construction of synthetic multi-cellular reaction-diffusion systems

We used synNotch receptors to design simple reaction-diffusion circuits (**Fig. 2A**). SynNotch receptors are mechanically activated juxtacrine receptors that can be used to engineer novel cell-cell signaling links (*20*). Based on the native Notch receptor, synNotch receptors undergo intramembrane cleavage when they bind their cognate ligand on a neighboring cell, releasing their transcription factor intracellular domain, allowing it to enter the nucleus and induce transcription of user-specified genes controlled by the synNotch responsive promoter. Diverse extracellular recognition domains and intracellular transcriptional domains can be used to program synthetic cell-cell regulatory links.

**Fig. 2.**
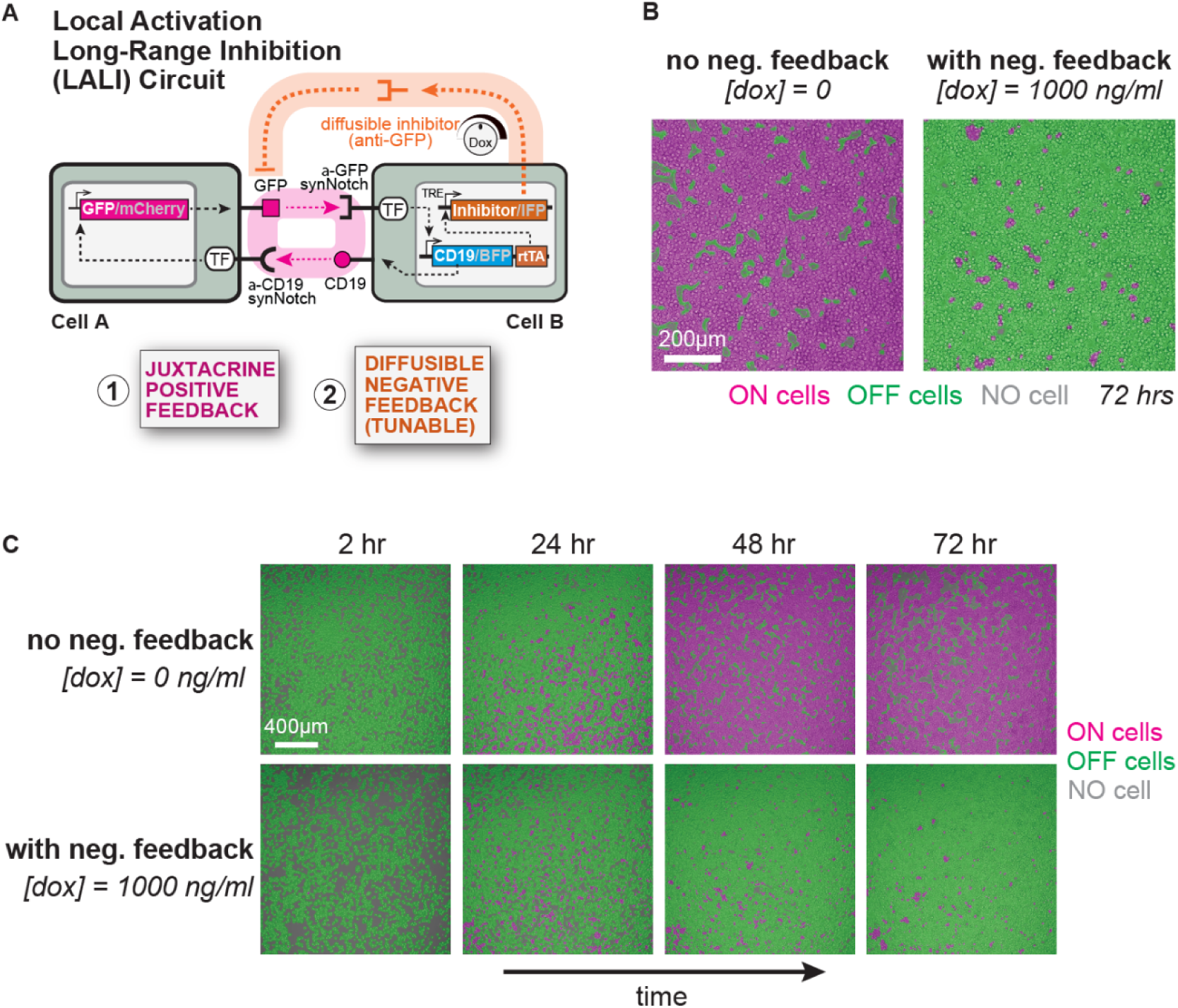
Modest pattern formation with core synthetic reaction diffusion circuits. (A) Synthetic LALI circuit design. Cell A expresses anti-CD19 synNotch that induces GFP_ligand_ and mCherry_reporter_. Cell B expresses anti-GFP synNotch that induces CD19_ligand_, BFP_reporter_ and rtTA which drives secretion of GFP_inhibitor_ and expression of IFP_reporter_. The induction level of GFP_inhibitor_ and IFP_reporter_ is tunable with Dox. See **Fig. S1** for more information on cell construction. (B) Synthetic pattern formation with 0 or 1000 ng/mL Dox. In the absence of Dox, the synthetic circuit functions as a positive feedback circuit, leading to spontaneous activation of nearly all cells. In the presence of 1000 ng/mL Dox, only a small population of cells was activated with strong negative feedback, forming a modest pattern with sparse tiny clusters. (C) Time course images of synthetic pattern formation by synthetic LALI circuit with 0 or 1000 ng/mL Dox. 20000 cells of each Cell A and Cell B were cocultured and imaged every 2hr for 72hr with variable Dox concentrations. Images at all time points were processed to define ON, OFF, and NO cell regions. See **Movie S1** for time-lapse videos.

Because synNotch signaling requires juxtacrine activation, it is inherently very short range. Thus, we can implement a short-range positive feedback loop by inducing expression of a synNotch ligand downstream from synNotch activation. Because expression of synNotch ligands and receptors on the same cell can lead to cis-inhibition, we decided to implement this kind of positive feedback loop using a two-cell mutual activation system: cell A has a synNotch receptor that detects the surface antigen CD19 and induces the expression of surface GFP, while cell B has a synNotch that detects surface GFP and induces expression of CD19 (**Fig. 2A**). Thus, activation of cell A would drive activation of cell B, and vice versa (two-cell autoactivation). In addition, in each cell, activation of its synNotch receptor drives the expression of a fluorescent reporter (either mCherry or BFP) as a marker of activation state. With this two-cell system, we can evaluate pattern formation by homogeneously mixing the two cell populations, plating in 2D, allowing them to interact, and observing the composite pattern of ON (either mCherry+ or BFP+) vs OFF (mCherry- or BFP-) cells (**Fig. 1C**).

We also could easily design a long-range negative feedback loop by having one of the synNotch receptors also drive expression of a diffusible inhibitor (**Fig. 2A**). In this case, we used a soluble anti-GFP nanobody as an inhibitor, as it competitively blocks binding of the GFP ligand to the anti-GFP synNotch receptor (*18*). Importantly, because this inhibitor is diffusible, it can signal at a longer range than the juxtracrine restricted synNotch receptor. To tune the ratio of positive and negative feedback, we drove expression of the diffusible inhibitor via a cascade in which the anti-GFP synNotch drives expression of rtTA, a doxycycline (Dox)-inducible transcriptional activator, which in turn induces expression of the inhibitor. In this way, there is no negative feedback in the absence of Dox, but negative feedback can be tunably increased, in the same cells, simply by adding increasing levels of Dox.

More concretely, using L929 mouse fibroblasts, we constructed the following two cell types (**Fig. 2A and Fig. S1**);

***Cell A:* anti-CD19 synNotch** → **[GFP_ligand_ + mCherry_reporter_]**

***Cell B:* anti-GFP LaG17 synNotch** → **[CD19_ligand_ + rtTA + BFP_reporter_] ; pTRE** → **[GFP _inhibitor_ + IFP_reporter_]**

The GFP_ligand_ and CD19_ligand_ are fusion proteins of GFP or CD19 extracellular domains fused to the PDGFR transmembrane domain, thus allowing the ligand juxtacrine presentation to the appropriate partner cell (*20*). The diffusible GFP inhibitor is a fusion protein of LaG16 and LaG2 anti-GFP nanobodies (*21*) that binds GFP and prevents its activation of anti-GFP LaG17 synNotch (*18*). We co-cultured 20,000 cells of each cell type in a non-treated 96-well plate and analyzed the emergence of spatial patterns of activated (mCherry+ or BFP+) and inactive (mCherry- or BFP-) cells over 72 hrs (**Fig. 1C and Fig. S2**).

### Core synthetic reaction-diffusion circuit yields only modest pattern formation

We first examined the dynamics of the synthetic reaction-diffusion circuit in the absence of Dox (i.e. no negative feedback), a situation in which there is only juxtacrine autoactivation. As expected, under this condition nearly all cells (both Cell A and Cell B) became uniformly activated within 72 hrs, spread across the entire plate surface (**Fig. 2B**, left panel, **Fig. 2C**, upper column, and **Movie S1**, left panel). This “all ON” behavior was expected, as the circuit with only positive feedback should lead to the propagation of activation throughout the entire population of cells.

We then examined what happened when we introduced long-range negative feedback in combination with juxtacrine positive feedback (core LALI reaction diffusion circuit). We co-cultured the cells (A and B) in the presence of Dox, which induces the long-range negative feedback link. Under high Dox conditions (1000 ng/ml Dox), which corresponds to maximum negative feedback strength, only a small fraction of cells became activated. Moreover, under these conditions the cells only self-organized into fairly modest patterns, showing formation of small, poorly defined spots, each made up of only a few cells (**Fig. 2B**, right panel, **Fig. 2C**, bottom column, and **Movie S1**, right panel).

### Coupling of induced cell adhesion to reaction-diffusion circuit improves pattern formation

Given the relatively poor pattern formation generated by our synthetic LALI circuit, we hypothesized that the circuit might require further fine tuning and optimization of parameters. We postulated, however, that there may be alternative ways to design LALI circuits that might enhance their robustness – their ability to show pattern formation across a wider range of parameters. Recent studies in model developmental systems have shown that biochemical morphogen signaling often operates in conjunction with cellular mechanical regulation to produce more robust spatial tissue patterns (*22–26*). For example, during hair follicle development, reaction diffusion based epithelial prepatterns are followed by aggregation of mesenchymal cells to form hair follicle spots, suggesting that integration of biochemical signaling and fate changes with mechanical sorting processes could be a mechanism to enhance robustness of multicellular patterning (*27–30*).

Based on this principle, we hypothesized that inducing aggregation of activated cells could enhance juxtacrine positive feedback and promoting more robust pattern formation. L929 cells (used here) have low background of cell-cell adhesion, and thus we could modulate their mechanical organization by inducing the expression of adhesion molecules such as Cadherins (*31*). To test this hypothesis, we engineered both Cell A and Cell B to induce expression of the cell adhesion protein P-cadherin (**Fig. 3A**), in conjuction with induction of the positive feedback synNotch ligands:

**Fig. 3.**
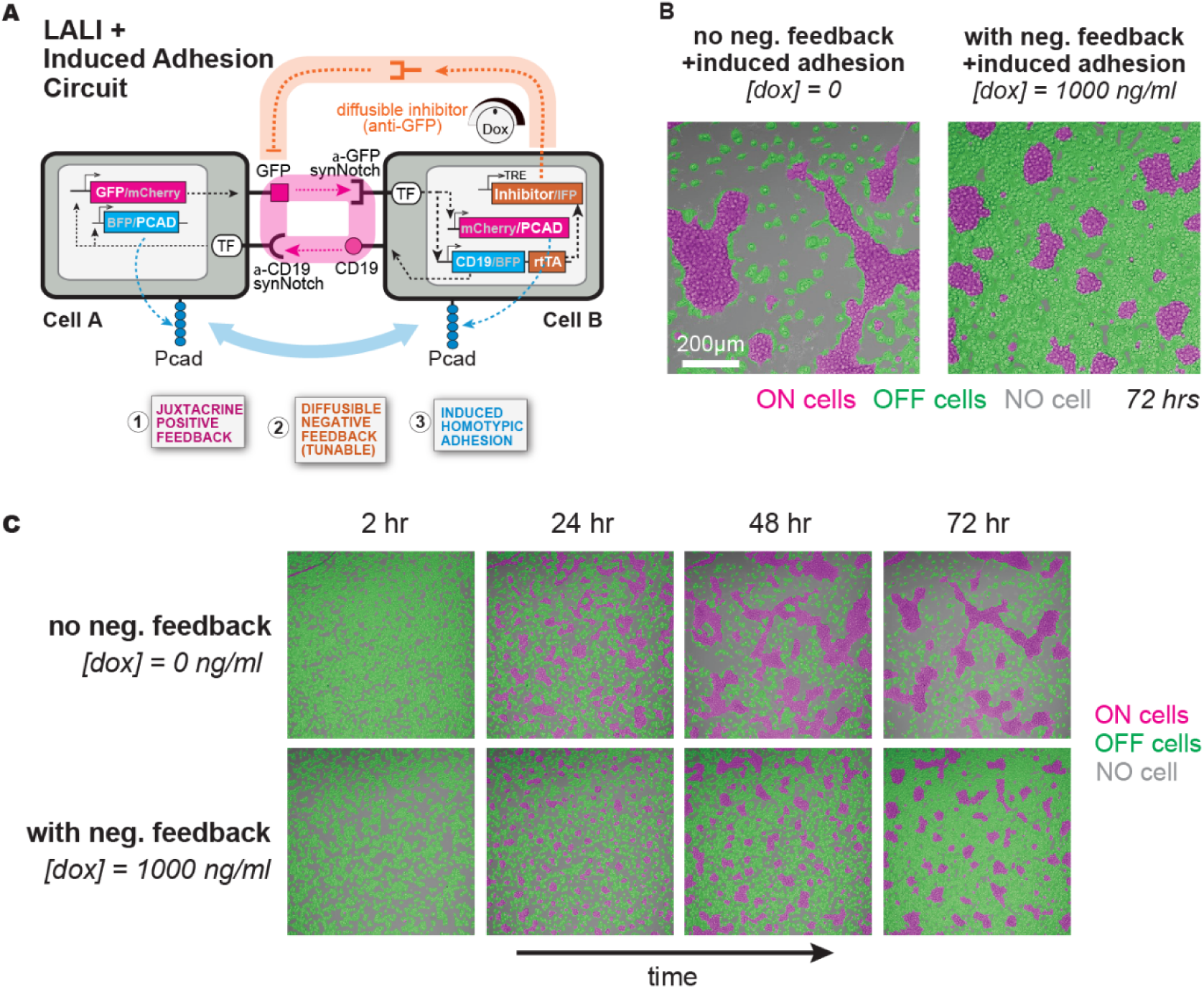
Adding induced adhesion to reaction diffusion circuit yields improved pattern formation. (A) Synthetic circuit design coupling LALI with induced cell adhesion. Cell A-iPcad expresses anti-CD19 synNotch that induces P-cadherin and BFP_reporter_ in addition to GFP_ligand_ and mCherry_reporter_. Cell B-iPcad expresses anti-GFP synNotch that induces P-cadherin and mCherry_reporter_ along with the original signaling components in Cell B. The induction levels of GFP_inhibitor_ in cell B-iPcad remain tunable with Dox. See **Fig. S3** for more information on cell construction. (B) Synthetic pattern formation with 0 or 1000 ng/mL Dox in adhesion-coupled circuits. In the absence of Dox, the synthetic circuit combining positive feedback and induced adhesion triggered widespread activation and formed large aggregates of most cells. In the presence of 1000 ng/mL Dox, diffusible negative feedback generated distinct spot-like clusters of ON cells, surrounded by OFF cells. (C) Time course images of synthetic pattern formation by synthetic LALI circuit with induced adhesion. 20000 cells of each Cell A-iPcad and Cell B-iPcad were cocultured and imaged every 2 hr for 72hr with variable Dox concentrations. Images at all time points were processed to define ON, OFF, and NO cell regions. See **Movie S2** for time-lapse videos.

*Cell A-iPcad:* **anti-CD19 synNotch** → **(GFP_ligand_ + mCherry_reporter_ and P-cadherin + BFP_reporter_)**

*Cell B-iPcad:* **anti-GFP LaG17 synNotch** → **(CD19_ligand_ + rtTA + BFP_reporter_ and P-cadherin + mCherry_reporter_) ; TRE** → **GFP_inhibitor_ + IFP_reporter_]**

Induction of P-cadherin expression in Cell A-iPcad and Cell B-iPcad was monitored using BFP and mCherry reporters, respectively (**Fig. S3A**). We confirmed that both cell types induced the expression of matched levels of P-cadherin upon activation, allowing the two cells type to remain well mixed within aggregates during co-culture experiments (**Fig. S3B**). Importantly, Dox-dependent IFP reporter induction in Cell B-iPcad was comparable to that in the original Cell B, indicating that the additional circuit engineering did not alter the induction levels of the diffusible GFP inhibitor that mediates long-range negative feedback (**Fig. S1** and **Fig. S3A**).

When Cell A-iPcad and Cell B-iPcad were co-cultured without Dox (no negative feedback), nearly all cells became activated and formed large aggregates due to induced P-cadherin expression (**Fig. 3B**, left panel, **Fig. 3C**, upper column, and **Movie S2**, left panel). The spaces between the activated cell aggregates were largely empty (no cells, rather than OFF cells). To then test the coupling of synthetic reaction diffusion circuit with induced adhesion, we co-cultured the same cells in the presence of 1000 ng/ml Dox, which turned on the diffusible negative feedback loop. Here, in contrast to the circuit without adhesion (**Fig. 2**), we observed very well defined and clear pattern formation with much larger spots of ON cells spaced within the predominantly OFF cells throughout the culture field (**Fig. 3B**, right panel, **Fig. 3C**, bottom column, and **Movie S2**, right panel). Thus, adding induced adhesion to the core LALI circuit led to stronger pattern formation.

### Exploring the range of positive/negative feedback ratios that lead to pattern formation

To better understand why the LALI circuit combined with induced cell adhesion led to significantly improved pattern formation, we more systematically scanned circuit behavior across different negative feedback strengths by titrating the concentration of Dox added to the system (**Fig. 4 and Fig. S4**). In the basic LALI synthetic reaction diffusion circuit (without induced adhesion), titrating the negative feedback strength led to a threshold like bifurcation – below 10 ng/mL Dox nearly all cells in the field switched ON, while above 50 ng/mL Dox, nearly all cells in the field were OFF (**Fig. 4A**). Only at the Dox concentration of 20 ng/mL did the circuit lead to roughly equivalent populations of ON/OFF cells. In this narrow transition range, the cells formed a more labyrinthine rather than spot pattern. At higher Dox concentrations, only a small proportion of cells turned ON, and these formed minimally sized scattered clusters, each made up of only a handful of cells. These results indicate that while the simple synthetic reaction diffusion circuit is capable of some degree of pattern formation, there is a limited range of parameters in which the positive and negative feedback strengths are properly balanced. Thus, the circuit lacks robustness in forming well-defined patterns.

**Fig. 4.**
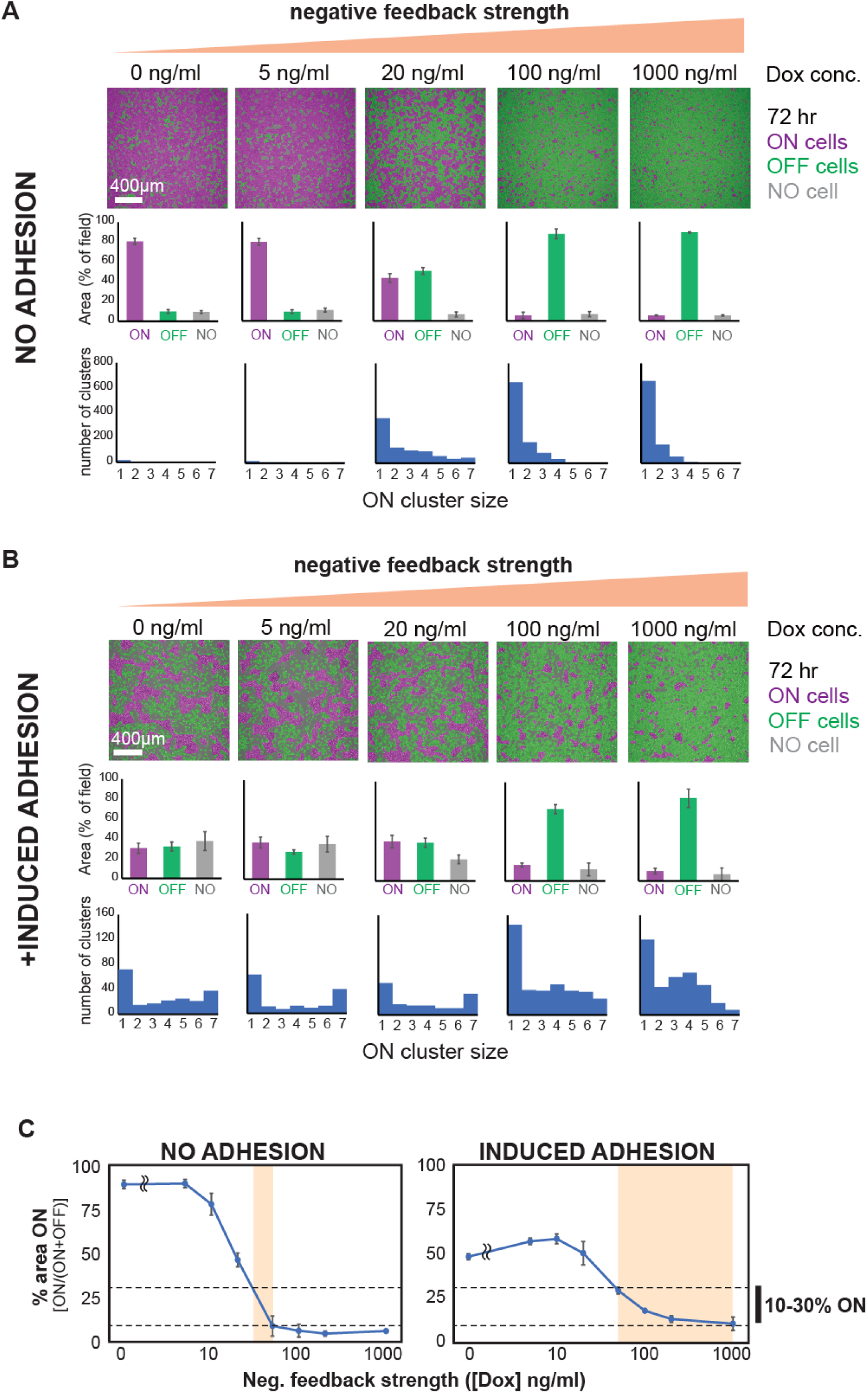
Combining induced adhesion to reaction diffusion signaling circuit leads to bigger range of negative feedback strength that yields well formed patterns. (A) Quantification of pattern formation in the original circuit. 20000 cells of each cell A and cell B were cocultured in the presence of variable Dox concentrations. The images were processed as described in **Fig. S2** to define ON, OFF, and NO cell regions, followed by quantification of each region area. The experiment was performed in triplicate and error bars show ±SD. Using the same data set, the sizes of ON cell clusters were quantified. The size distribution of ON cell clusters are shown by histograms with varying size categories in the x-axis (1: 100∼500 μm², 2: 500∼1000 μm², 3: 1000∼2000 μm², 4: 2000∼4000 μm², 5: 4000∼8000 μm², 6: 8000∼16000 μm², 7: >16000 μm²). The small clusters less than 100 μm² (< single cell) were cut off from the histograms. (B) Quantification of pattern formation in the adhesion-coupled circuit. 20000 cells of each Cell A-iPcad and Cell B-iPcad were cocultured with variable Dox concentrations, followed by image acquisition and processing as described in the methods. The area ratio of ON, OFF, and NO cell regions and the size distribution of ON cell clusters are shown as described in (A). (C) Comparison of ON cell fractions (% of ON cell area in ON and OFF cell area) across various negative feedback strengths. The synthetic LALI circuit showed a very narrow range of negative feedback strength for generating 10-30% ON cells, whereas, adhesion-coupled LALI circuits maintained 10-30% ON cell fractions robustly across a broader range of negative feedback strengths.

We performed a similar negative feedback strength scan on the LALI circuit with induced adhesion (**Fig. 4B**). With no negative feedback, nearly all cells turned ON and formed an aggregated labyrinthine network, with very few cells found in the intervening spaces. This pattern was expected, as the positive feedback alone should lead to all cells expressing high cadherin and strong aggregation of all cells. As we increased Dox concentration, the fraction of ON cells did decrease, but more gradually, such that significant populations of both ON and OFF cells were observed over a broader range of negative feedback strengths. This more balanced bifurcation of ON/OFF cells suggested that the coupling of induce cell adhesion made the circuit more robust over a wider range of negative/positive feedback strengths. Through a wider range of negative feedback strengths, a significant ON cell populations persisted and formed distinct spot-like clusters surrounded by a field of OFF cells (**Fig. 4B**). In contrast to the circuit without induced adhesion, ON-cell populations were maintained across a wider range of Dox concentrations, and cluster sizes remained large (most clusters were >10-cells and >1000 μm^2 in size) compared with the clusters in the circuit without induced adhesion (**Fig. 4A-B**). For example, when comparing the two circuits, we saw that the negative feedback strength range (Dox concentrations) that results in 10-30% ON cells is very narrow for the circuit without induced adhesion, but much wider for the circuit with induced adhesion (**Fig. 4C**). We hypothesize that induced cell adhesion effectively converts transient stochastic activation into more persistent and self-reinforcing activated spatial domains (i.e. adhesion will locally promote more short-range positive feedback). As a result, synthetic pattern formation would be less sensitive to the precise negative feedback strength. Thus, coupling reaction diffusion signaling with induced cell adhesion appears to substantially enhances the robustness of multicellular pattern formation.

### Agent-based computational framework for theoretical analysis of synthetic cellular reaction diffusion circuits

To theoretically investigate how coupling reaction-diffusion signaling circuits with induced cell adhesion enhances the robustness of multicellular patterning, we developed an agent-based model that incorporates our cell-cell communication circuits with regulated mechanical properties of cells (**Fig. 5A**). The model includes two cell types, Cell A and Cell B, with synNotch circuit designs corresponding to those implemented in our experiments (**Fig. 2**). Cells mutually activate target gene expression upon contact, and activated Cell B also secretes a GFP inhibitor that diffuses via random walk and suppresses anti-GFP synNotch activation. To model cellular adhesion interactions, we defined a distance-dependent force function between contacting cells (**Fig. 5B**). Cells without adhesion experience no attractive force, whereas cells with adhesion experience attractive forces as a function of distance. When cells get too close, repulsive forces arise. This distance-dependent force curve was quantitatively validated for cadherin-mediated cell-cell interactions (*32, 33*). In circuits with induced cell adhesion, the strength of attractive force increases with cadherin expression level at a fixed distance.

**Fig. 5.**
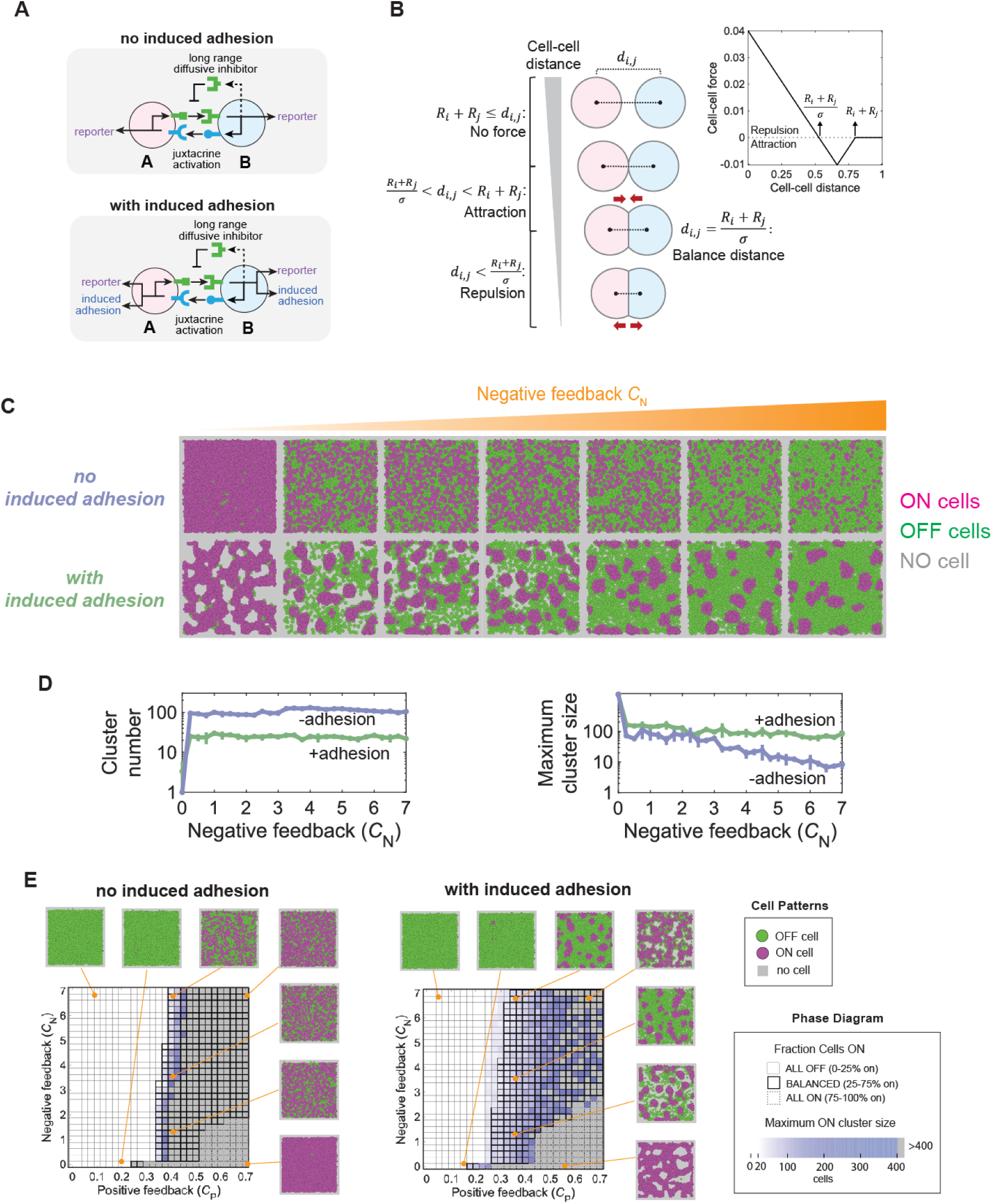
Computational modeling shows increased robustness to feedback strengths when reaction-diffusion circuit is combined with induced cell adhesion. (A) Schematic illustration of synthetic two cell types (Cell A and Cell B) with juxtacrine activation and long range diffusive inhibitor as implemented experimentally, shown for conditions with (bottom row) and without (top row) induced adhesion. (B) Distance-dependent force function between cells, where cells exert volume-driven repulsion at short distances and adhesion-driven attraction at moderate distances, provided the center-to-center distance remains smaller than the sum of their radii. (C) Final patterns of ON (magenta) and OFF (green) cells (gray: no cell) across varying negative feedback strengths, shown for conditions with (bottom) and without (top) induced adhesion. (D) Quantification of cluster number and maximum cluster size across varying negative feedback strength. (E) Phase diagram across a broad parameter space spanning positive and negative feedback strengths, shown for conditions with (right panel; with a wide parameter regime supporting spot pattern) and without (left panel; with a narrow parameter regime supporting spot pattern) induced adhesion, alongside ON cell fraction and maximum cluster size quantified and representative spatial patterns displayed.

At the beginning of simulations, equal numbers of Cell A and Cell B were randomly distributed on a square grid under periodic boundary conditions to eliminate edge effects (**Fig. S5**). Starting with a basal activation level, cells dynamically updated their activation levels and spatial positions through signaling feedback, induced adhesion, and stochastic movement by random walk. While some cells showed dynamic status changes between active and inactive during simulations, simulations were run over a duration long enough for the system to reach a steady state (**Fig. S5**). At the end of the simulation procedure, cells were classified as ON or OFF based on whether their activation level was over a defined threshold, enabling quantitative analysis of clustering behavior. All mathematical formulas and parameter values are detailed in Materials and Methods.

First, we tested whether our core synthetic reaction diffusion circuits can theoretically generate tissue patterns under idealized conditions (without cell motility and division) (**Fig. S6**). Under these conditions, spot or stripe-like patterns emerged at a very specific strength of negative feedback, suggesting that our synthetic circuits can form patterns but only under finely-tuned feedback parameters. We then incorporated realistic variability of cellular properties such as motility and stochasticity into the simulations. In agreement with the experimental results (**Fig. 2**), in the absence of negative feedback, the circuit alone led to nearly uniform activation, whereas increasing negative feedback reduced the fraction of activated cells (**Fig. 5C**, upper column). However, activated cells remained highly dispersed and failed to yield stable ON clusters. These results indicate that, although the core synthetic LALI circuit is capable of pattern formation, in principle, it is insufficient to robustly generate spatially organized multicellular patterns under realistic conditions.

In contrast, simulations incorporating both the synthetic reaction-diffusion circuit and induced cell adhesion produced stable spot-like patterns of ON cells surrounded by OFF cells across a wide range of inhibitory feedback strengths (**Fig. 5C**, bottom column). By stabilizing spatial positions of activated cells, adhesion seems to reduce the effective impact of stochastic cell movement and signal fluctuations. Induced adhesion likely acts as an additional positive feedback component, as close cell clustering mediated by adhesion will reinforce the short-range juxtacrine positive feedback signaling. Compared with the synthetic reaction diffusion alone, induced cell adhesion maintained both the number and size of ON-cell clusters across varying negative feedback strengths (**Fig. 5D**). Thus, the agent-based model recapitulated the experimental results and demonstrates that coupling reaction diffusion signaling with cell adhesion substantially enhances patterning robustness.

### Coupling reaction-diffusion signaling with physical cell adhesion expands parameter space that generates strong pattern formation

To systematically assess pattern-forming capacity of our synthetic reaction diffusion circuits under diverse signaling conditions, we used our agent based computational model to explore a wide parameter space spanning positive and negative feedback strengths, with or without induced cell adhesion (**Fig. 5E**). Phase diagrams were generated to characterize cell fate bifurcation (ON-vs-OFF ratio) and cluster size distributions under variable conditions of positive and negative feedback strength. Here we considered good pattern forming circuits to be those that yield relatively balanced ratios of ON-vs-OFF cells, and to generate medium sized ON cluster sizes (defined spots).

The core synthetic reaction-diffusion LALI circuit exhibited a very narrow slice parameter space that permits pattern formation, suggesting that robust multicellular patterning is difficult to achieve through reaction diffusion signaling alone in this system (**Fig. 5E**, left panel). In contrast, coupling the circuit with induced cell adhesion dramatically expanded the area of parameter space that supports stable spot-like patterns (**Fig. 5E**, right panel).

To examine whether the effect of signaling-adhesion coupling during reaction diffusion patterning can be generalized beyond our specific circuit implementation, we applied the same modeling framework to a more general cases of LALI reaction diffusion systems (**Fig. S7A**). For example, rather than using a two-cell type positive feedback loop, we also modeled a single cell type circuit, in which one cell express a receptor that induces the expression of both a short-range activator and a long-range inhibitor of the receptor. By systematically scanning feedback parameters, we generated phase diagrams describing the capacity of cell fate bifurcation and clustering behavior. We observed the same trend in a single cell configuration of a LALI circuit that we observed for the two-cell configuration. Incorporation of induced cadherin-mediated cell adhesion consistently enables better pattern formation across a broader range of system parameters (**Fig. S7B**). Together, these results demonstrate that integrating reaction–diffusion signaling with mechanical cell adhesion provides a general strategy for achieving spontaneous and robust multicellular pattern formation.

## Discussion

In this study, we combined experimental and theoretical analyses of synthetic cellular reaction diffusion circuits to investigate minimal genetic programs required for robust multicellular patterning. Our initial experimental synthetic reaction diffusion circuit exhibited spontaneous activation, but proportional cell fate bifurcation and pattern formation occurred only within a very narrow range of negative feedback strengths. At low negative feedback strengths, the vast majority of cells turned ON, while at higher negative feedback strengths, the majority turned OFF, limiting the range in which ON-OFF patterns could be observed. Moreover, even in these ranges, the activated cells were individually distributed without forming clearly defined spatial patterns. These results indicate that our simple LALI synthetic signaling circuit alone lacks robustness, and highlights the utility of this synthetic system to explore factors that can make reaction diffusion patterning more robust. Leveraging the modularity of synNotch engineered cell regulatory circuits, we tested the impact of integrating induced cell adhesion into our synthetic reaction diffusion circuits. In silico exploration additionally provided a framework to explore a much larger design space than accessible through experiments alone. Rational integration of both experimental and in silico approaches therefore offers a powerful strategy to evaluate and construct robust multicellular systems.

A common challenge in designing and building complex biological systems, like those capable of self-organized patterning, is achieving robust behaviors. Evolved or designed systems, especially in a process as important as development, must be highly resistant to noise and fluctuations (*34–37*). Real world cell systems contain diverse stochastic fluctuations that could affect or disrupt reaction reaction-diffusion processes, including cell proliferation, cell motility, and cell-to-cell variability in ligand and receptor expression levels as well as target gene induction levels. Fine-tuning all network parameters and controlling such stochastic fluctuations in mammalian cells is inherently challenging.

Here we have demonstrated that coupling mechanical cell aggregation/sorting to reaction-diffusion signaling provides a key mechanism that can increase the robustness of self-organizing pattern forming cellular circuits. For LALI reaction-diffusion circuits to generate patterns on their own, factors such as negative/positive feedback ratios and diffusion rate ratios must be finely balanced to yield heterogenous patterns of ON and OFF cells. We postulate that coupled adhesion and mechanical sorting can cooperate with these morphogen driven circuits by more tightly aggregating ON cells and reinforcing their local positive feedback activation (**Fig. 6A**). This model is consistent with the observation that the coupled circuit yields ON-cell aggregate spots with a physically protruding, raised 3D morphology (**Fig. 6B**).

**Fig. 6.**
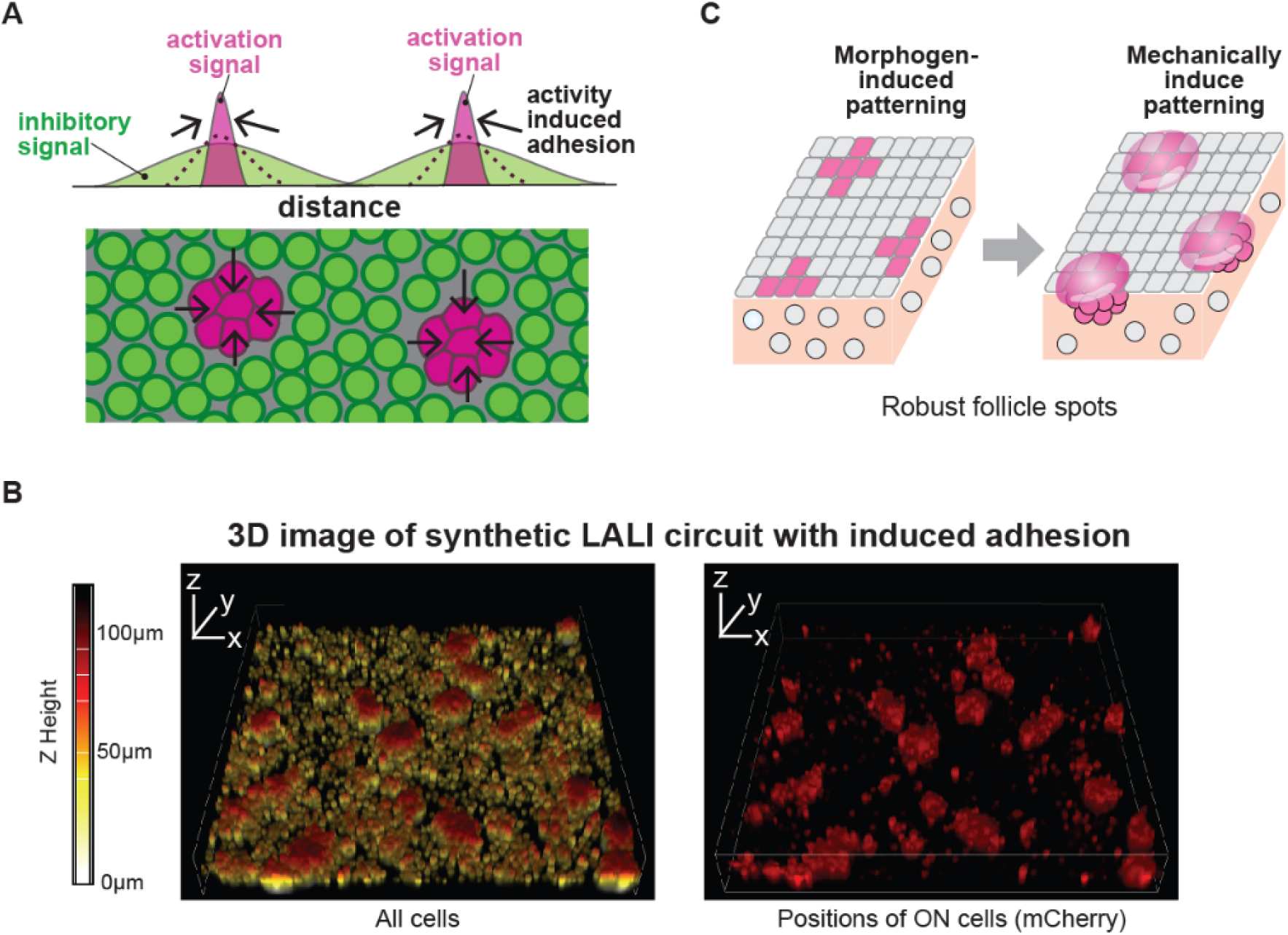
Cooperative integration of signaling with mechanical changes leads to robust spot pattern formation. (A) Cooperation of biochemical LALI circuit and mechanical cell aggregates. LALI circuit spontaneously generates local activation, while activity-induced adhesion restricts the activation signal into local spots, leading to stabilization of activated clusters and enhancing robustness. (B) 3D imaging of synthetic patterns generated by synthetic LALI circuit coupled with induced adhesion. Cell A-iPcad and Cell B-iPcad cells were stained with CellTrace Far red dye and cocultured in the presence of 1000 ng/mL Dox for 72 hr. The red hot scale showed z-axis heights of synthetic patterns. (C) Design principles for robust spot formation through the interplay of morphogen-induced patterning and mechanically-inducing patterning.

This principle of cooperation between morphogen-mediated signaling and mechanical adhesion changes is found in natural developmental processes such as skin follicle pattern formation (*27*) and zebrafish spinal chord patterning (*24*). Thus, this principle likely extends beyond our specific system (**Fig. 6C**). During animal evolution, patterning mechanisms that couple morphogen signaling and adhesion likely emerged, and their coupling may have conferred robustness to developmental processes, enabling the evolution of increasingly complex tissue architectures. Thus, coupling of morphogen-based positional instructions with mechanical rearrangements driven by forces like differential adhesion may be fairly ubiquitous across many developmental processes (*38, 39*).

We show here how generative de novo design of synthetic self-organizing cell circuits provides a way to systematically dissect and probe minimally sufficient circuit features robust pattern formation. Our design of synthetic cellular reaction diffusion circuits does not rely on native morphogens, but rather employs contact-dependent synNotch signaling as a local activator and a secretion-based GFP inhibitor as a long-range inhibitor. These contact-based activating signals can function as slowly-diffusing activators (compared to the diffusible inhibitor), enabling the construction of LALI-type circuits. Of course every circuit, whether evolved or designed, has its individual idiosyncracies, but our more complete theoretical analysis exploring alternative circuit implementations (e.g. single cell positive feedback, diffusible activators/inihibitors; Fig S8), indicates that the principle outlined here – that coupling of adhesion based sorting with LALI circuits generates more robust pattern formation – is very general.

Although the current study focuses primarily on key features such as negative/positive feedback strength and coupling of adhesion, there are other circuit parameters that could be examined in a similar manner using a growing set of tools with which to systematically reconstitute self-organizing systems. For example, there are emerging tools to tune the diffusion rate/distance of signaling molecules (*40, 41*). These and other new synthetic morphogenic tools will provide powerful approaches to both dissect developmental principles and to potentially engineer tissues with greater precision and robustness.

## Materials and Methods

### Plasmid construct design for synthetic cellular reaction diffusion circuit

We built synthetic cellular reaction diffusion circuits based on synthetic Notch receptors (*20*). Anti-GFP LaG17 synNotch receptor (Addgene, #79127) was constructed by fusing human CD8a signal sequence, myc tag, anti-GFP nanobody LaG17 (*21*) for extracellular recognition domain, mouse Notch1 (NM_008714) transmembrane regulatory region (Ile1427 to Arg 1752), and Gal4-VP64 for intracellular transcription factor domain. Anti-CD19 synNotch receptor was constructed by fusing human CD8a signal sequence, myc tag, anti-CD19 scFv for extracellular recognition domain and same transmembrane and intracellular domains of Anti-GFP LaG17 synNotch receptor. GFP_ligand_ was constructed by fusing mouse IgK signal sequence, EGFP, and PDGFR transmembrane domain (Addgene, #79129). CD19_ligand_ was constructed by fusing mouse IgK signal sequence, HA tag, human CD19 extracellular domain, and PDGFR transmembrane domain (Addgene, #133803). GFP_inhibitor_ was constructed by fusing Gaussia Luciferase signal sequence, myc tag, and GFP nanobody LaG16-LaG2(*21*). We assembled these synNotch ligands and receptors, fluorescent reporters (BFP, mCherry and IFP (IFP2.0 fused with nuclear localized signal)), puromycin resistant gene (puroR), GFP inhibitor, Neomycin resistant gene (NeoR), P2A sequence and mouse P-cadherin into a modified pHR’SIN:CSW vector with UAS, TRE3GS and PGK promoters to construct following plasmids that were used to build the Cell A, Cell B, Cell A-iPcad, and Cell B-iPcad. Cell A:

1. pHR_UAS→GFP_ligand_-P2A-mCherry_reporter__PGK→anti-CD19 synNotch-P2A-puroR

Cell B:

1. pHR_UAS→CD19_ligand_-P2A-rtTA-P2A-BFP_reporter__PGK→anti-GFP LaG17 synNotch-P2A-puroR
2. pHR_TRE3GS→GFP_inhibitor_-P2A-IFP_reporter__PGK→NeoR

Cell A-iPcad:

1. pHR_UAS→GFP_ligand_-P2A-mCherry_reporter__PGK→anti-CD19 synNotch-P2A-puroR
2. pHR_UAS→BFP_reporter_-P2A-P-cadherin_PGK→puroR

Cell B-iPcad:

1. pHR_UAS→CD19_ligand_-P2A-rtTA-P2A-BFP_reporter__PGK→anti-GFP LaG17 synNotch-P2A-puroR
2. pHR_TRE3GS→GFP_inhibitor_-P2A-IFP_reporter__PGK→NeoR
3. pHR_UAS→mCherry_reporter_-P2A-P-cadherin_PGK→puroR

#### Cell culture

Mouse fibroblast line L929 (RIKEN BRC, Cell Bank) was cultured in DMEM/10%FBS (DMEM (Nacalai tesque, #08458-16) containing 10% fetal bovine serum (Gibco) and 1x penicillin-streptomycin solution (Wako, #168-23191)) at 37 °C in humidified environment with 5% CO2. L929 cells were detached by 3-5 min incubation at 37 °C in TrypLE Express (Gibco, #12605028) from TC-treated culture dish. Cells were then spun down by 400xg 3min centrifuge and counted for plating. Human erythroleukemic cell line K562 (ATCC, #CCL-243) was maintained with DMEM/10%FBS in non-treat culture dish.

#### Lentiviral transduction

All engineered L929 cells in this study were stably transformed with lentiviral transduction. Lentiviral supernatant was prepared by transfecting the pHR’SIN:CSW vector and the packaging plasmids pCMVdR8.91 (*42*) and pMD2.G (Addgene #12259) into HEK293T cells using PEI MAX (Polysciences, #24765). 1.0x10^5^ L929 cells were infected with variable amounts of lentiviral supernatant in the presence of 10 μg/mL hexadimethrine bromide (Sigma-Aldrich, #H9268) in 12-well plates for 48 h.

##### Construction of synNotch-expressing cells (Cell A, Cell B, Cell A-iPcad, and Cell B-iPcad)

To construct Cell A, L929 cells were transduced with the plasmid described above by lentiviral infection, split into two wells of 12-well plate, and incubated overnight. To stimulate anti-CD19 synNotch, 8.0x10^5^ K562 cells expressing the CD19_ligand_ (K562/CD19) were added onto the infected cells and cocultured for 24hr (*20*). Following stimulation, the basal/induced expression levels of mCherry reporter were measured using a cell sorter (SH800S, SONY). Cells were then single-cell sorted into 96-well plates. We then waited for the growth of sorted clones for around 15 days and subsequently split into three 96-well plates. Following two days incubation, one plate was stimulated with 4.0 × 10⁴ K562/CD19 cells per well and cocultured overnight, while the remaining plates were left unstimulated. The 96-well plates with or without stimulation by K562/CD19 were analyzed with a cell analyzer (SA3800, SONY). The unstimulated 96-well plate was used as a control to measure background level of target gene expression. Based on the reporter fluorescence intensity with/without K562/CD19 stimulation, we selected clones for further experiments. The flow cytometry data were analyzed with FlowJo software.

To construct Cell B, L929 cells were first transduced with Plasmid 1 (pHR_UAS→CD19_ligand_-P2A-rtTA-P2A-BFP_reporter__PGK→anti-GFP LaG17 synNotch-P2A-puroR) using a procedure similar to that described for Cell A. At this time, we used K562 cells expressing the GFP_ligand_ (K562/GFP) to stimulate anti-GFP LaG17 synNotch. After establishing a clone expressing Plasmid 1, we lenti-virally introduced Plasmid 2 (pHR_TRE3GS→GFP_inhibitor_-P2A-IFP_reporter__PGK→NeoR) into the clone. Cells were then stimulated with K562/GFP in the presence of 1000 ng/ml Dox. The basal/induced expression levels of BFP and IFP reporters were measured, and cells were single-cell sorted into 96-well plates (SH800S, SONY). We screened clones based on the fluorescence intensity of reporters with/without K562/GFP stimulation. The Dox-dependent IFP reporter induction levels in activated Cell B were further quantified with a cell analyzer (CytoFLEX S, Beckman Coulter) in the presence of various concentrations of Dox indicated in **Fig. S1**.

To construct P-cadherin-inducible lines, Cell A-iPcad and Cell B-iPcad, we lenti-virally introduced the plasmids (pHR_UAS→BFP_reporter_-P2A-P-cadherin_PGK→puroR or pHR_UAS→mCherry_reporter_-P2A-P-cadherin_PGK→puroR) into Cell A or Cell B, respectively. Cells were then stimulated with K562/CD19 or K562/GFP to activate anti-CD19 synNotch or anti-GFP LaG17 synNotch. The basal/induced expression levels of BFP and mCherry reporters were measured, and cells were single-cel sorted into 96-well plates (SH800S, SONY). We screened clones based on the fluorescence intensity of reporters with/without K562/CD19 or K562/GFP stimulation. The Dox-dependent IFP reporter induction levels in activated Cell B-iPcad were quantified with a cell analyzer (CytoFLEX S, Beckman Coulter) in the presence of various concentrations of Dox to confirm the comparable induction.

#### Pattern formation assay

20,000 cells of two cell type (Cell A/Cell B or Cell A-iPcad/Cell B-iPcad) were mixed and plated in 200ul of DMEM/10%FBS in non-treat 96 well plate (Corning #3370). Subsequently, 50ul of DMEM/10%FBS containing Dox at variable concentrations were added at final concentrations indicated in the figures. The plate was then centrifuged at 100xg for 2 min to plate cells uniformly in the wells. We incubated the plate in the stage-top chamber incubation system (TOKAI HIT) equipped on Nikon AX-R confocal microscope for **Fig. 2**, **Fig. 3**, **Fig. 4B**, and **Fig. S4B**. Cells were maintained at 37 °C in a humidified atmosphere with 5% CO₂ by supplying water and CO₂ to the chamber. The multi-well time lapse imaging was performed using the Nikon AX-R confocal microscope equipped with a CFI Plan Apochromat Lambda D 10X objective (0.45 NA) (Nikon) and controlled by JOBS in NIS-Elements software (Nikon) to automatically capture the images at the center of each well with 10 μm-step three or five z-slices every 2 hr for 72 hr. The maximum intensity projection images were generated with z-slice images for further processing and analysis. For endpoint imaging in **Fig. 4A**, **Fig. S3B** and **Fig. S4A**, the plate was incubated in the CO₂ incubator for 72hr and imaged using the Nikon AX-R confocal microscope as described above. For 3D imaging in **Fig. 6B**, both Cell A-iPcad and Cell B-iPcad were stained with Cell Trace Far red dye (ThermoFisher #C34572) and cocultured in the presence of 1000 ng/mL Dox. The plate was incubated for 72 hr in the stage-top chamber incubation system (TOKAI HIT) equipped on Keyence BZ-X810. After incubation, 3 μm-step 39 z-slices were captured using the Nikon AX-R confocal microscope with a Plan Apochromat Lambda 20X objective. The images were processed with denoise.ai in NIS-Elements software (Nikon) to generate 3D images of Cell Trace Far red dye (all cells) and mCherry (ON cells) channels.

#### Image processing and analysis

We created masks for activated cells, inactive cells, and no cell space (ON cells, OFF cells, and NO cell) in the acquired images. We then quantified the area of ON/OFF/NO cell regions to calculate their ratio and created histograms of ON cell masks to show their size distribution. We processed 1024x1024 pixels images with following procedures using ImageJ macro.

ON cell mask for coculture experiments of Cell A and Cell B (Magenta):

To generate ON cell masks in **Fig.2**, **Fig. 4A**, **Fig. S4A** and **Fig. S5A**, images were processed as shown in **Fig. S2**. We first normalized the induced levels of mCherry and BFP reporters in Cell A and Cell B respectively. We quantified mCherry and BFP intensity induced by positive feedback circuit (with no Dox) and measured the intensity values that reached 20% saturation. We then run “setMinAndMax” to apply the range (from 0 to the measured values) to the histograms of mCherry and BFP. We merged the normalized mCherry and BFP images and binarized them to create the images showing cell activation status. To reduce the spatial resolution for further analysis, we run "Gaussian Blur" with radius = 4 pixels and binarized the processed images to define activated regions. We applied magenta and green pseudocolors to ON and other regions, respectively.

### No cell mask (Dark grey)

To define the area with NO cell, we processed bright field images. First, we run “Subtract Background” with rolling ball radius = 50 pixels and then run "Find Edges" and "Gaussian Blur" with radius = 4 pixels to find and fill the cell area. We binarized the processed images and converted no cell area to NO cell mask with dark grey pseudocolor.

### OFF cell mask (Green)

We defined the area other than ON cell mask and NO cell mask as OFF cell mask.

### ON cell mask for coculture experiments of Cell A-iPcad and Cell B-iPcad (Magenta)

To generate ON cell masks in **Fig. 3**, **Fig. 4B**, and **Fig. S4B**, we used mCherry reporter for both Cell A-iPcad and Cell B-iPcad, as mCherry expression is induced at comparable levels in both cell types upon activation by K562/CD19 or K562/GFP (**Fig. S3**). We first normalized mCherry intensity such that activated cells in coculture images of Cell A/Cell B or Cell A-iPcad/Cell B-iPcad exhibit the same saturation level. Because activated Cell A-iPcad and Cell B-iPcad formed highly compact and tall aggregates with no Dox, % of ON cell area of Cell A-iPcad/Cell B-iPcad could shrink compared to that of Cell A/Cell B. We therefore selected images at 10 ng/mL Dox for normalization where most cells were still activated but exhibited reduced compaction. To define the saturation threshold, we measured the saturation rate of merged images of normalized mCherry and BFP signals in the regions containing cells of the Cell A and Cell B coculture at 10 ng/mL Dox. The same saturation rate was then applied to the images of the Cell A-iPcad and Cell B-iPcad coculture at 10 ng/mL Dox by adjusting the maximum value of histogram range of mCherry. The same histogram range was applied as a saturation threshold to normalize all images at varying Dox concentrations in **Fig. 4B** and **Fig. S4B**. We then binarized the normalized images, and run "Gaussian Blur" with radius = 4 pixels, and binarized the processed images to define activated regions. We applied magenta and green colors to ON and other regions, respectively.

#### Agent-based model

The agent-based model – that describes each microscopic entity (*e.g.*, a cell or molecule) as an individual agent – is often used for simulating the interactions and motions within a multicellular system. Despite coarse-graining microscopic entities to single points for computational efficiency, this model still uncovers the fundamental principles of multicellular patterning in embryogenesis, organogenesis, *etc.* (*43, 44*). Here, we utilize this approach for its exceptional computational efficiency *via* dimensional reduction, enabling modeling spatiotemporal dynamics of thousands of entities (*incl.*, cells and molecules).

### 1) Intercellular force

The mathematical formulation for pairwise intercellular repulsive (volume repelling) and attractive (cell adhesion) forces is adopted from (*32, 33, 45*), which was proven capable of inferring heterogeneous adhesion among neighboring cells during embryonic development.

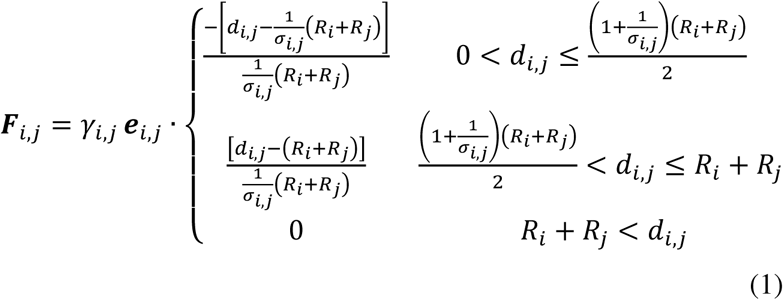

where i_i,j_ denotes the force imposed on Cell i from Cell j; i_i,j_ denotes the unit vector pointing from Cell j to Cell i; d_i,j_ denotes the distance between Cell *i* and Cell j ; R_i_ and R_j_ denote the radius of Cell i and Cell j ; i_i,j_ denotes the repelling strength between Cell i and Cell j and is positive; a_i,j_ denotes the adhesion strength between Cell i and Cell j and is no smaller than 1; 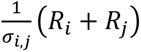 denotes the balance distance between Cell i and Cell j where the interacting force is zero; when 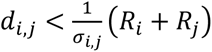, the force is repulsive; when 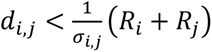, the force is attractive. A schematic diagram of distance-force relationship is shown in **Fig. 5B**.

### 2) Signaling feedback in a two-cell-type circuit

The synthetic multicellular system consisting of cell type A and B is involved with diffusion-based (long-range) negative signaling feedback and contact-based (short-range) positive signaling feedback. The gene expression of a B-type cell leads to a proportional release of diffusive molecules (C_N_S_i_ new negative-feedback molecules per time step; S_i_ denotes the gene expression of Cell *i*) that are initially produced on its periphery, proceed random walking, and exist for a lifespan of T_M_:

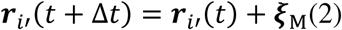

where the Molecule i′ is subjected to Gaussian noise ξ_M_ in all orthogonal spatial directions, and the gene expression of a B-type cell will be inhibited by the diffusive molecule number (n_N_) within its cell periphery.

Prior to the inhibition, the gene expression of either a A-type or B-type cell is activated by the contact with the other:

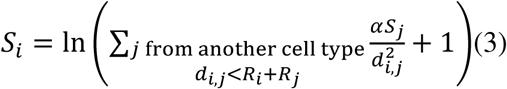

then, for cell type A, S_i_ is further multiplied by C_P2_ to denote the strength of contact-based activation; for cell type B, S is further multiplied by 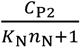 to denote the strength of diffusion-based inhibition on top of contact-based activation (K_N_ denotes the inhibition response and is no smaller than 0. S_i_ is thresholded to a lower limit S_min_, considering the intrinsically stochastic and basal gene expression).

### 3) Signaling feedback for a one-cell-type circuit

The comparative multicellular system consisting only one cell type is involved with diffusion-based (long-range with much faster random working on a molecule) negative signaling feedback and diffusion-based (short-range with much slower random working on a molecule) positive signaling feedback. The gene expression leads to a proportional release of two different kinds of diffusive molecules (C_P1_S_i_ new positive-feedback molecules and C_N_S_i_ new negative-feedback molecules per time step; S_i_ denotes the gene expression of Cell *i*) that are initially produced on its periphery, proceed random walking, and exist for a lifespan of T_M_:

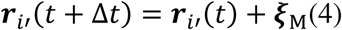

where the Molecule i′ is subjected to Gaussian noise ξ_M_ in all orthogonal spatial directions, and the gene expression will be inhibited by the diffusive molecule number (n_N_) within its cell periphery.

Prior to the inhibition, the gene expression is activated by the diffusive molecule number (n_P1_) within its cell periphery:

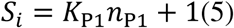

then, S_i_ is further multiplied by 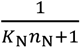 to denote the strength of diffusion-based inhibition on top of diffusion-based activation (K_N_ denotes the inhibition response and is no smaller than 0. S_i_ is thresholded to a lower limit S_min_, considering the intrinsically stochastic and basal gene expression).

### 4) Induced cell adhesion

The gene expression of two neighboring cells mediates their cell adhesion strength by:

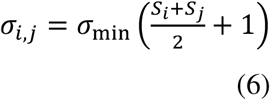

where a_min_ is the basal adhesion strength before additional gene regulation; a_i,j_ is thresholded to an upper limit a_max_, considering that cells cannot approach each other very closely due to volume repelling, even when gene expression and adhesive protein levels overshoot.

### 5) Cell movement under signaling and mechanics

Each cell moves based on all intercellular forces imposed on it, along with Gaussian noise ξ_C_ in an overdamped environment and within periodic boundaries:

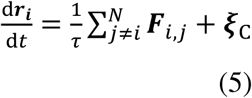

Here, if the right-hand term exceeds the substrate friction 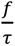, a motion friction force of magnitude 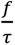 and opposite direction is added; otherwise, the entire right-hand term is nullified by static friction.

### 6) Pattern initiation, simulation, and quantification

A total of N A-type cells and N B-type cells in a two-cell-type circuit or N identical cells in a one-cell-type circuit are randomly arranged in a square grid of width and height L, with a gene expression level of S_min_. Starting from this initial condition, the simulation proceeds for a total duration t_max_ with a time step Δt. At the final time point, ON and OFF cells are binarized by applying a gene expression threshold S_T_. Isolated clusters of ON cells are then determined based on their neighborhood connectivity, following the approach described in previous 2D adhesive cell pattern study (*46*). Periodic boundary conditions are validated by confirming that a homogeneous initial cell pattern remains intact when random cellular and molecular processes are disabled and the parameters γ, β_min_, f, r, and N are varied.

## Acknowledgement

We thank Dr. Kosuke Mizuno for discussion and testing on image analysis. We also thank former and current members of the Toda laboratory and the Lim laboratory for discussion and assistance.

## Author contributions

S.T. developed and planned research; carried out design, construction, and testing of multicellular patterning circuits; and wrote the manuscript. G.G. planned research; carried out design, construction, and testing of agent-based modeling and simulation; and wrote the manuscript. M.T. and H.M. helped testing of multicellular patterning circuits. W.A.L. developed and oversaw research; and wrote and edited the manuscript.

## Data and materials availability

All data are available in the main manuscript and supplementary materials. All plasmids developed in this study will be deposited at Addgene, where they will be publicly available.

---

**Table S1.**
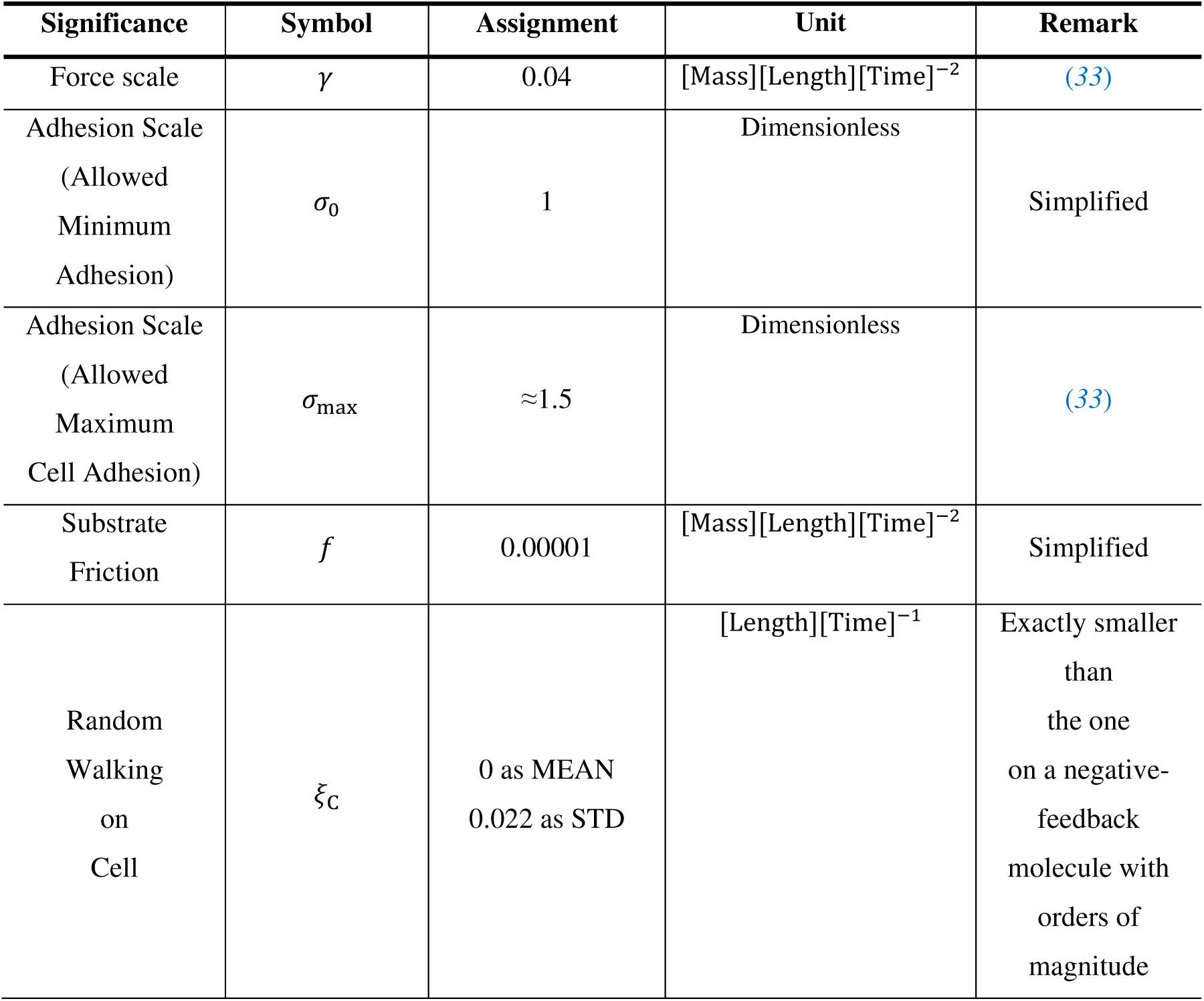
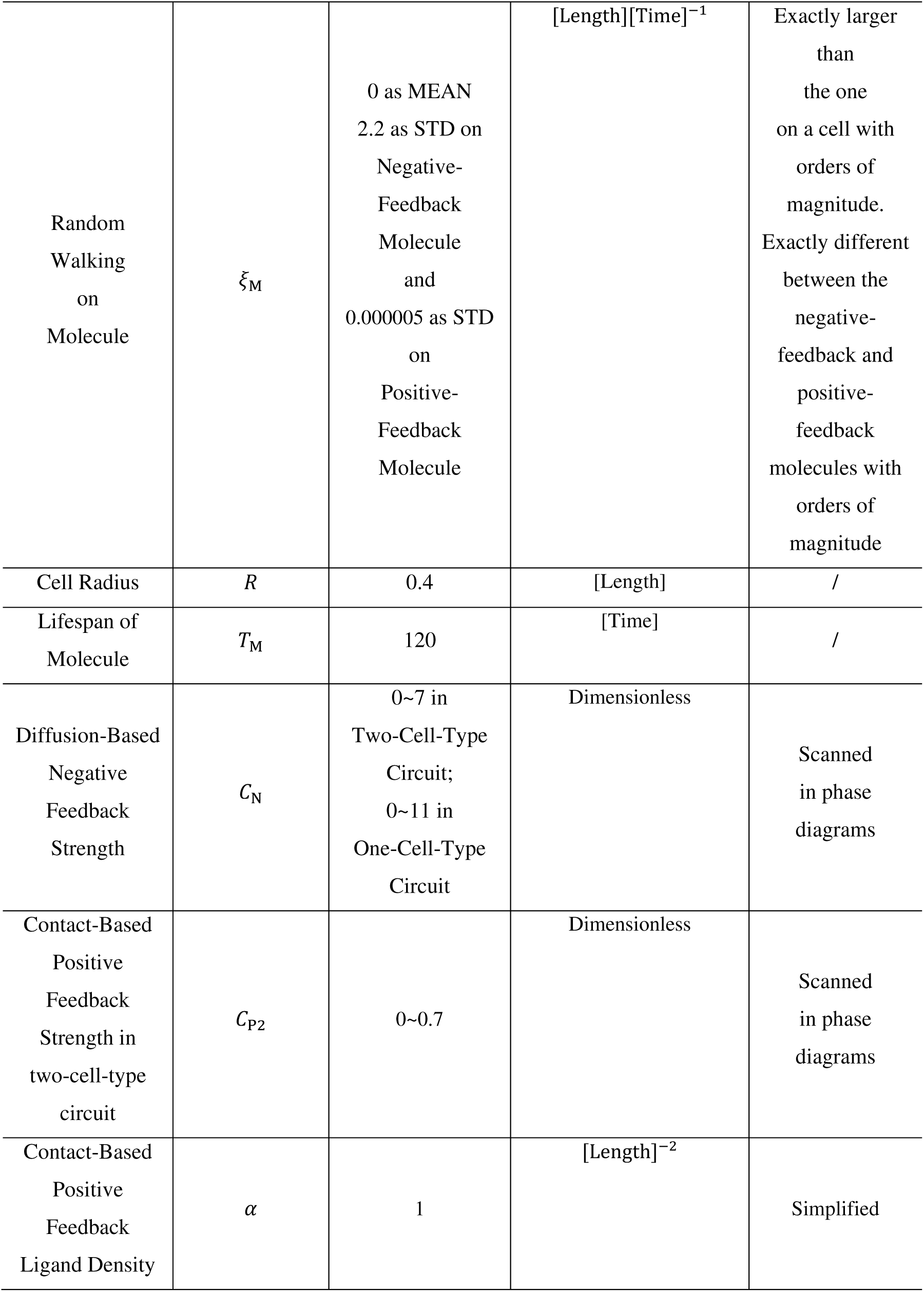
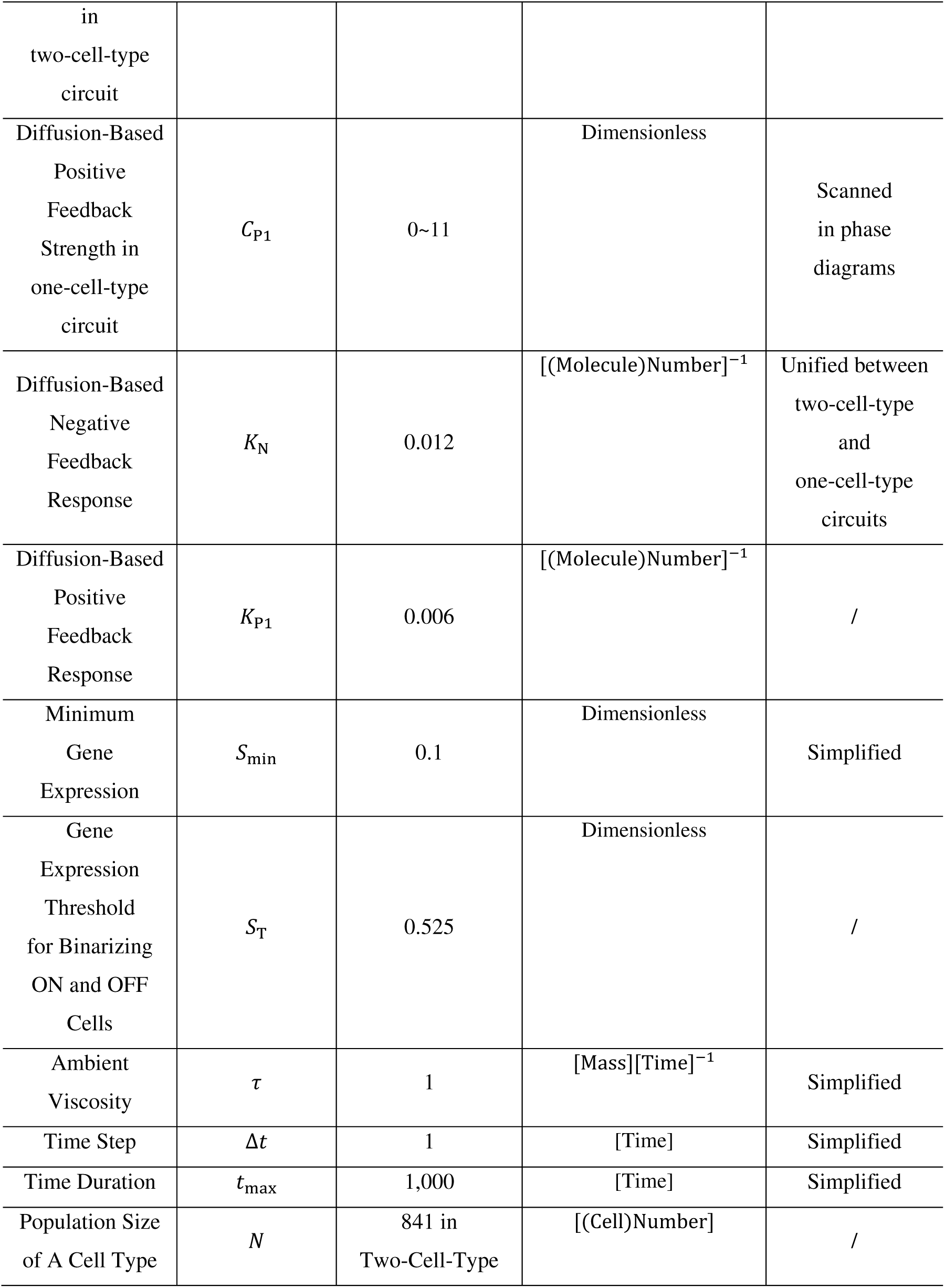
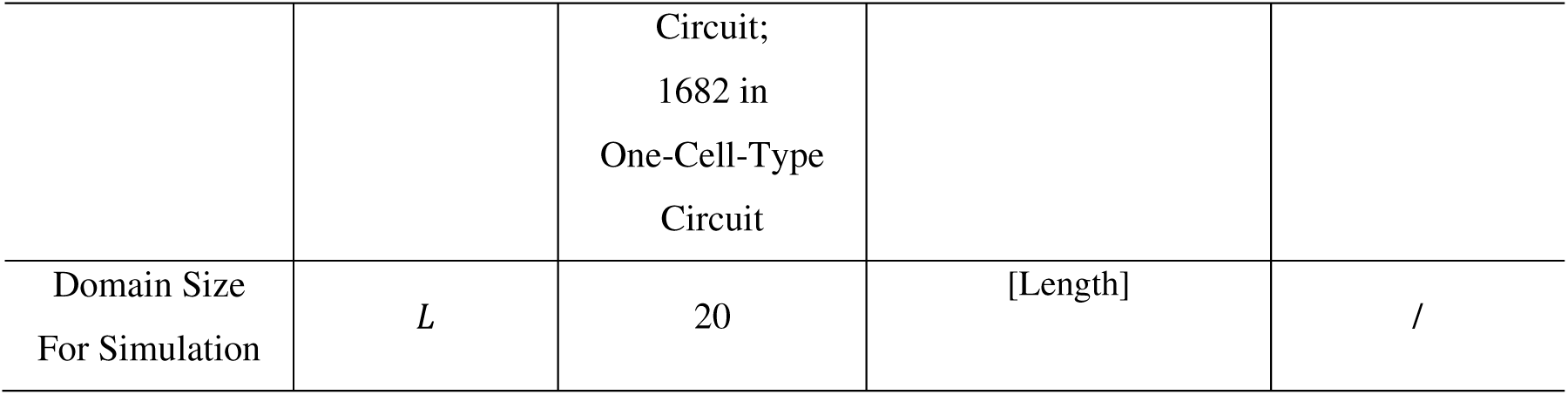
Model parameter information for the default setting.

**Fig. S1.**
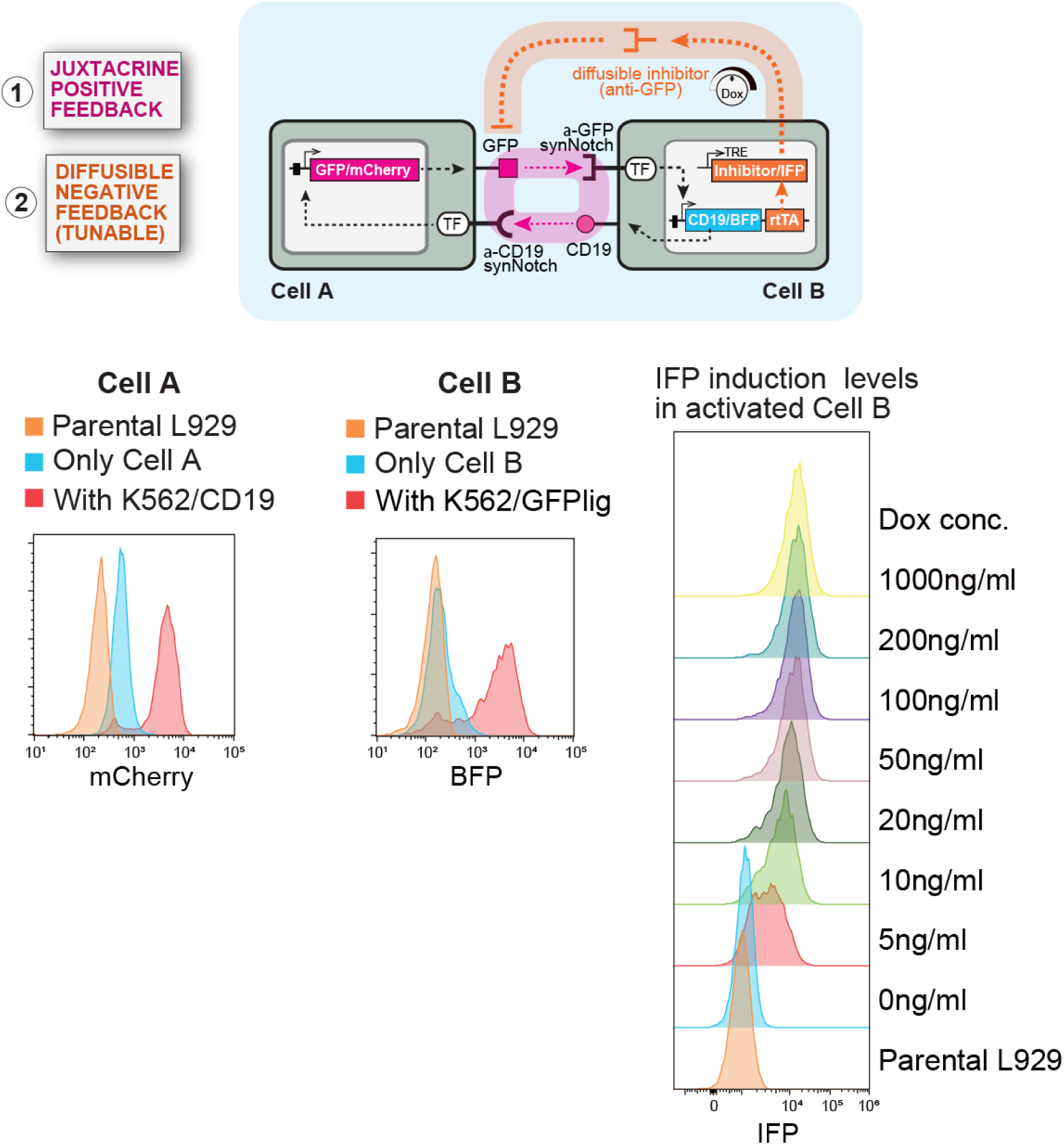
Construction of Cell A and Cell B that form juxtacrine positive feedback circuit with tunable and diffusible negative feedback. Cell A: We engineered L929 cells to express anti-CD19 synNotch that induces membrane-tethered GFP_ligand_ and mCherry_reporter_. The expression level of mCherry_reporter_ was measured with or without stimulation by CD19_ligand_-expressing K562 cells (K562/CD19) using flow cytometry. Cell B: We engineered L929 cells to express anti-GFP LaG17 synNotch that induces CD19_ligand_, BFP_reporter_ and rtTA. The expression level of BFP_reporter_ was measured with or without synNotch stimulation by GFP_ligand_ -expressing K562 cells (K562/GFP) using flow cytometry. Cell B was further engineered to induce the expression of GFP_inhibitor_ and IFP_reporter_ by rtTA in the presence of Dox. The induction levels of IFP_reporter_ in activated Cell B were tunable with variable Dox concentrations.

**Fig. S2.**
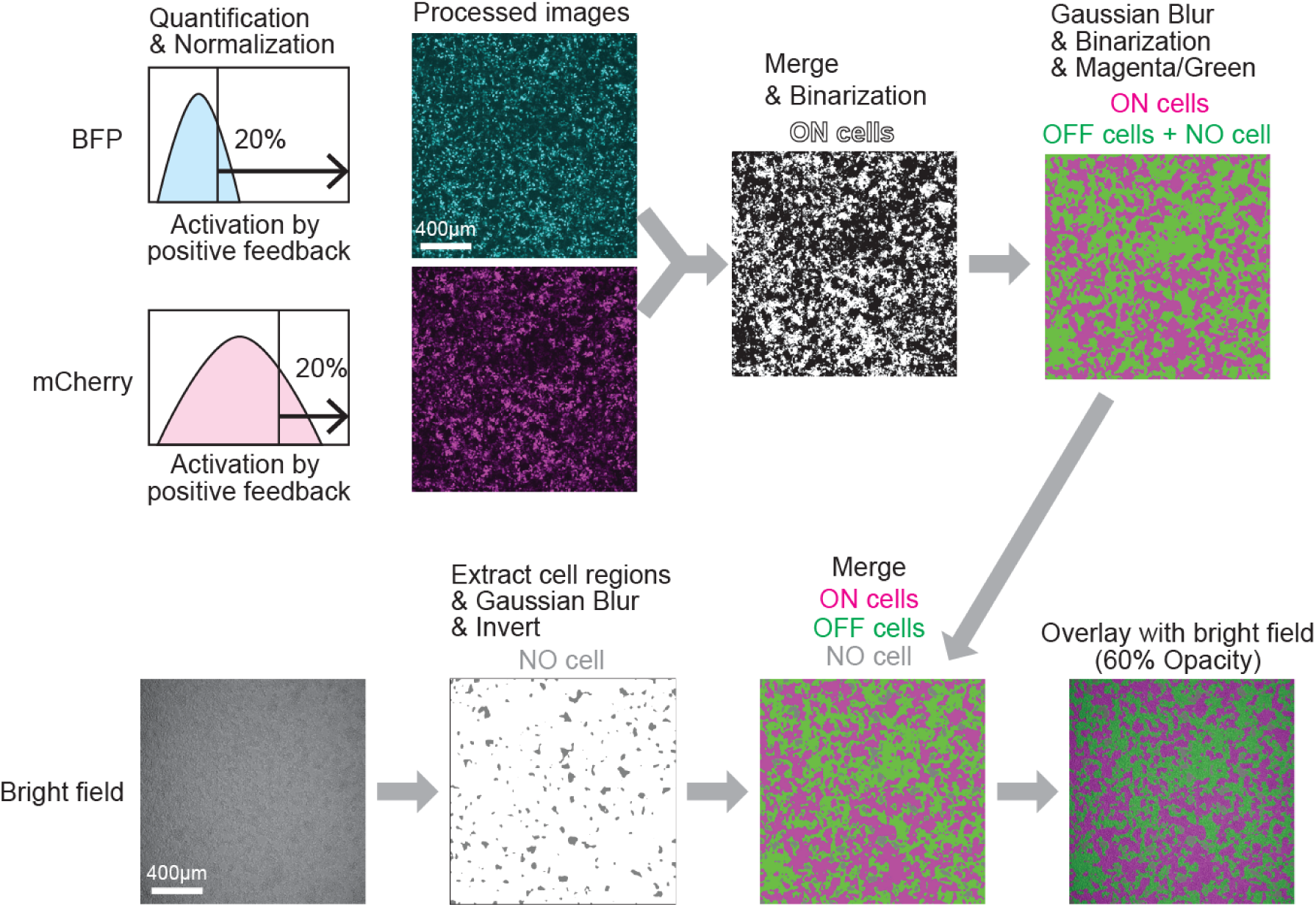
Image processing pipeline to create masks for ON, OFF, NO cell regions. Workflow for image processing to define the area of activated cells (ON cells), inactive cells (OFF cells) and no cell space (NO cell) based on fluorescence and bright-field images. First, we quantified the mCherry and BFP reporters induced in Cell A and Cell B respectively when activated by positive feedback circuit with no Dox. We measured the intensity value that reached 20% saturation in each channel and normalized BFP and mCherry intensity to create merged images and define ON cells with binarization. For further analysis, we run Gaussian Blur for spatial smoothing and binarized the processed images to define the regions of ON and OFF cells that include NO cell regions. To define NO cell regions, we extracted cell regions from bright field images using Find Edges and Gaussian Blur and inverted them. Finally, we overlaid bright field images to binarized images of ON, OFF and NO cells.

**Fig. S3.**
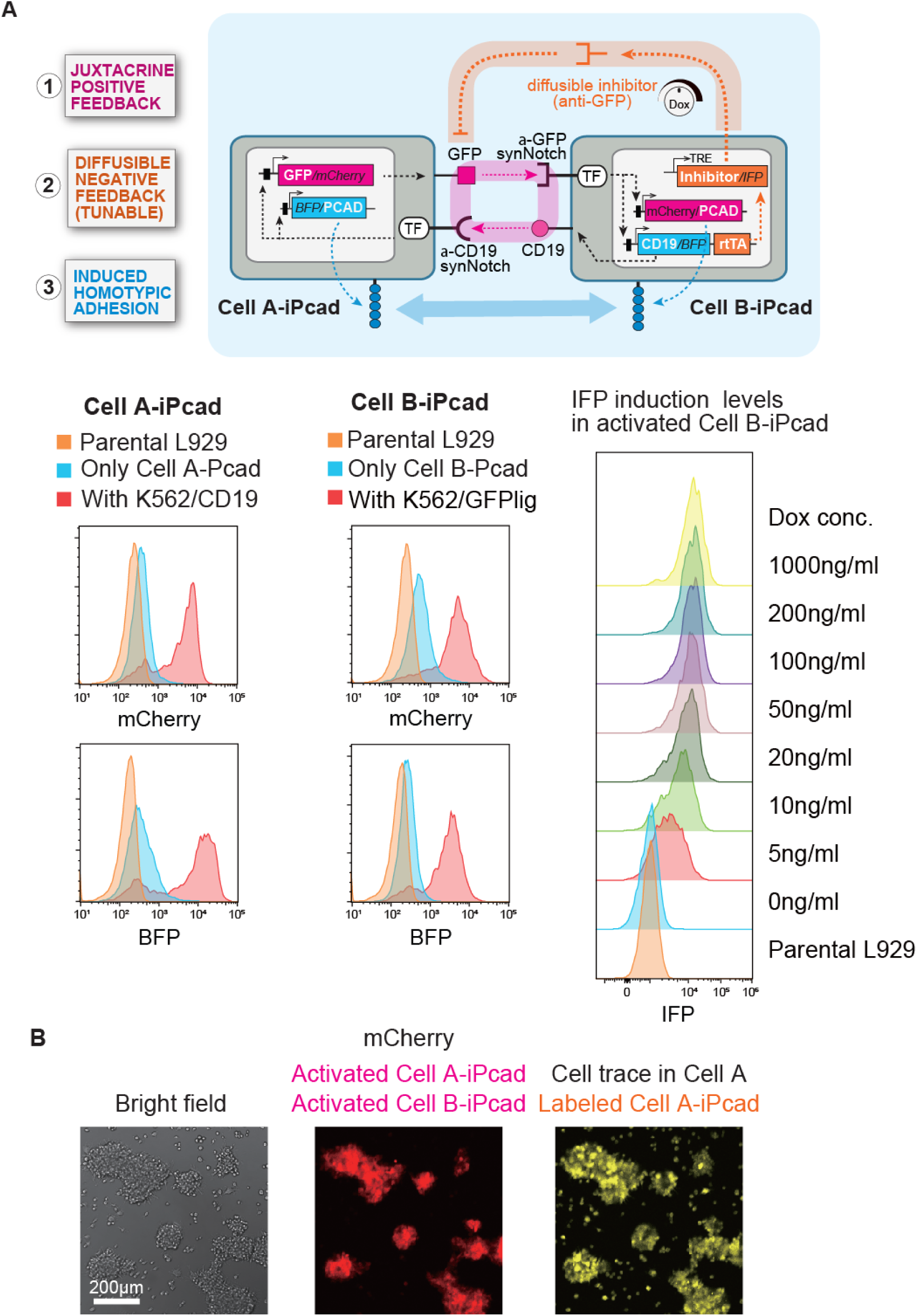
Construction of Cell A and Cell B with induced P-cadherin expression. (A) Cell A-iPcad: Cell A was further engineered to induce P-cadherin and BFP_reporter_ in the downstream of anti-CD19 synNotch. The expression level of both mCherry_reporter_ and BFP_reporter_ were measured with or without stimulation by K562/CD19 using flow cytometry. Cell B-iPcad: Cell B was further engineered to induce P-cadherin with mCherry_reporter_ in the downstream of anti-GFP LaG17 synNotch. The expression level of both BFP_reporter_ and mCherry_reporter_ were measured with or without synNotch stimulation by K562/GFP using flow cytometry. The induction levels of IFP_reporter_ in Cell B-iPcad show similar tunability to Cell B in the presence of variable Dox concentrations. (B) Comparable adhesion induction in Cell A-iPcad cells and Cell B-iPcad. 20000 cells of each Cell A-iPcad and Cell B-iPcad were cocultured in the absence of Dox, leading to activation through positive feedback and formation of large cell aggregates. By labeling Cell A-iPcad with CellTrace Far red dye, well-mixed distributions of Cell A-iPcad cells in the aggregates were visualized, suggesting that both cell types induced similar levels of P-cadherin expression when activated.

**Fig. S4.**
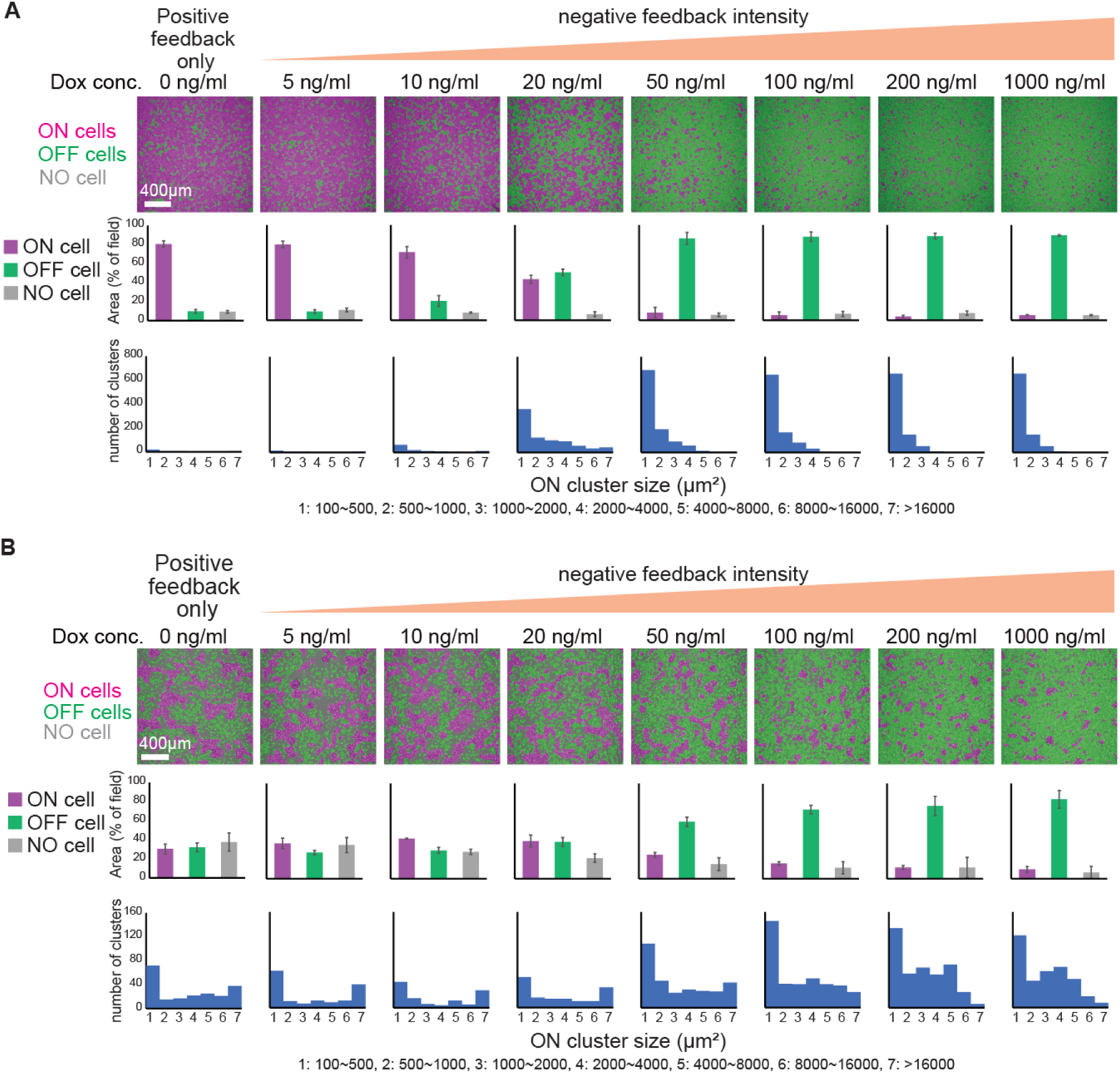
Complete data sets of synthetic pattern formation used for quantification in Fig. 4C. (A) A complete data set of synthetic pattern formation by coculturing Cell A and Cell B with variable Dox concentrations used for quantification. (B) A complete data set of synthetic pattern formation by coculturing Cell A-iPcad and Cell B-iPcad with variable Dox concentrations used for quantification.

**Fig. S5.**
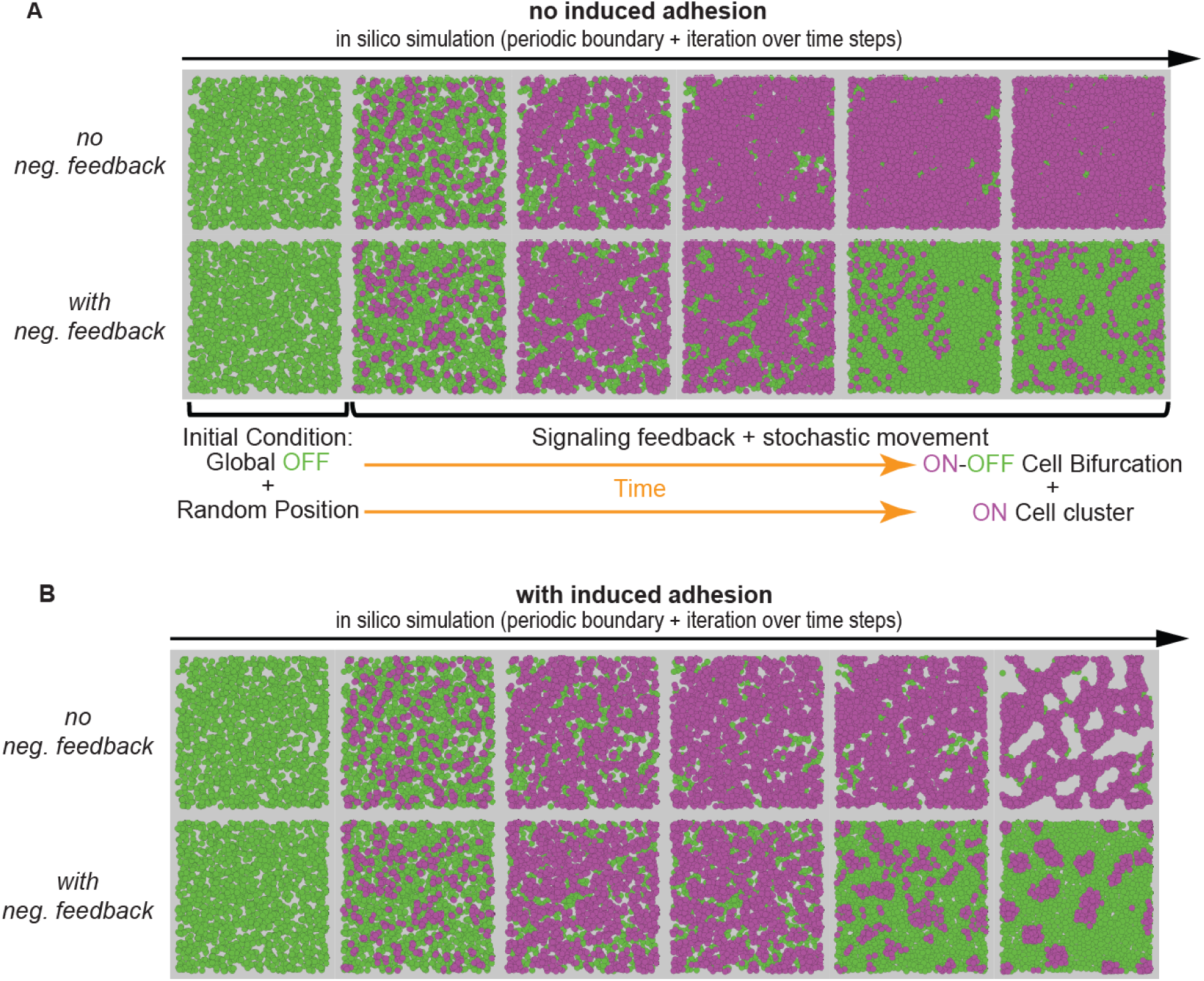
Computational modeling of synthetic LALI circuits. (A) Intermediate patterns of ON (magenta) and OFF (green) cells (gray: no cell) from initial to final moments when there is negative feedback (bottom row) or not (top row), shown for condition without induced adhesion. (B) Intermediate patterns from initial to final moments when there is negative feedback (bottom row) or not (top row), shown for condition with induced adhesion. Simulation settings are indicated along with the time-lapse series.

**Fig. S6.**
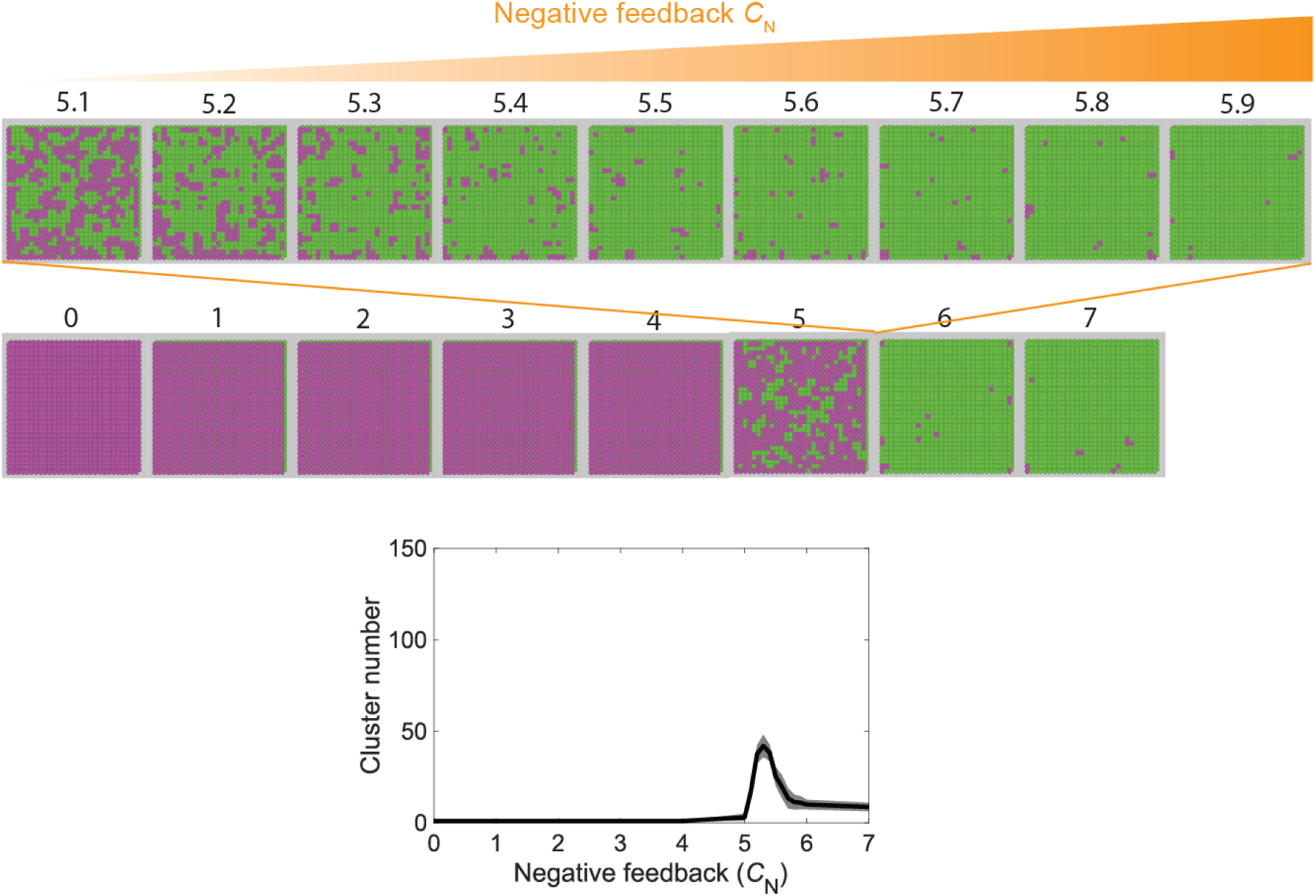
Synthetic reaction diffusion circuits (without induced adhesion) can induce pattern formation within a very narrow range of negative feedback at ideal conditions. Final patterns of ON (magenta) and OFF (green) cells (gray: no cell) across varying negative feedback strengths, alongside cluster number quantified beneath.

**Fig. S7.**
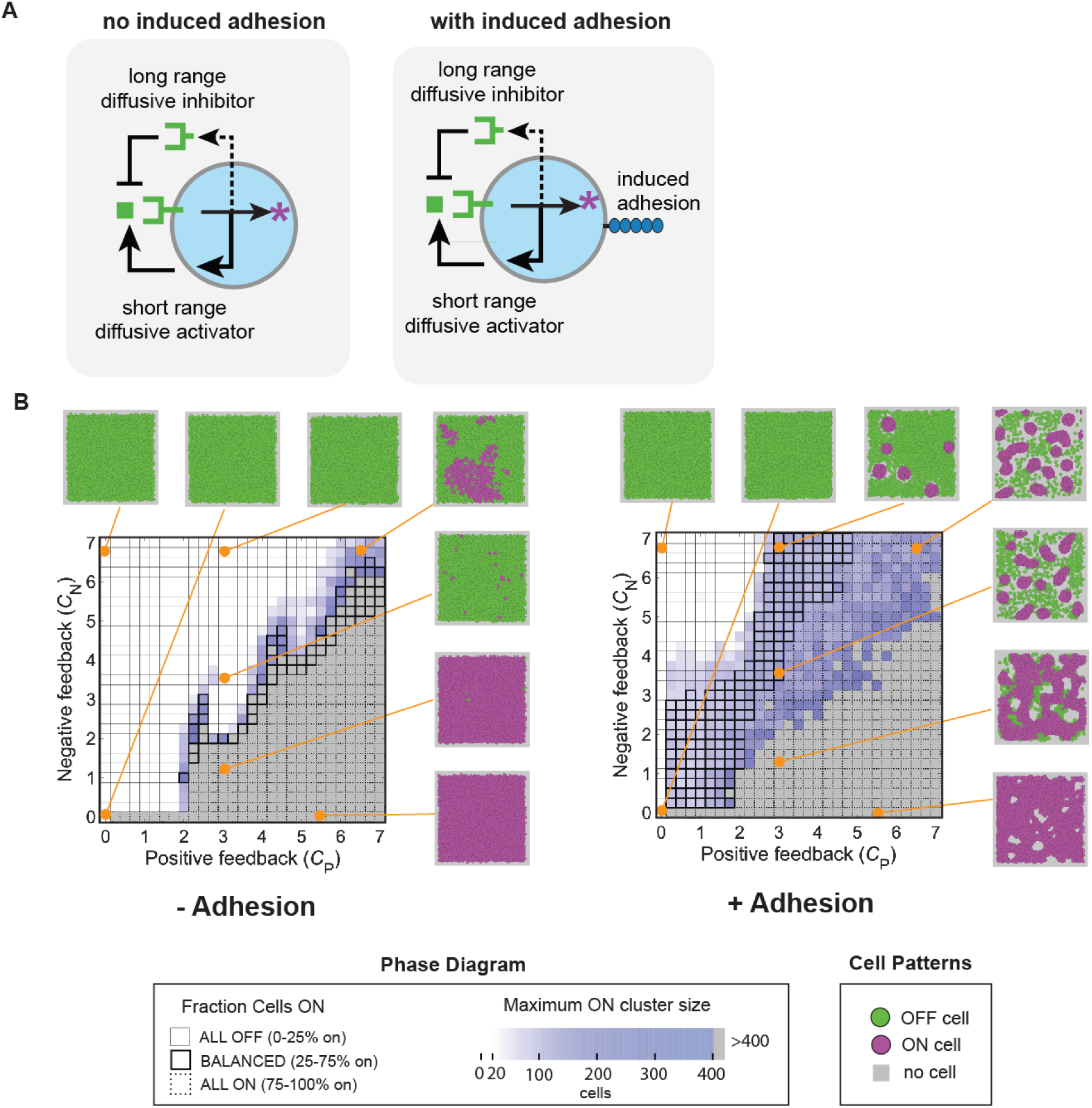
Generalizing robust reaction-diffusion patterning coupled with cell adhesion in single cell circuit and diffusible activator and inhibitor. (A) Schematic illustration of single cell type with short range diffusive activator and long range diffusive inhibitor, shown for conditions with (right panel) and without (left panel) induced adhesion. (B) Phase diagram across a broad parameter space spanning positive and negative feedback strengths, shown for conditions with (right panel; with a wide parameter regime supporting spot pattern) and without (left panel; with a narrow parameter regime supporting spot pattern) induced adhesion, alongside ON cell fraction and maximum cluster size quantified and representative spatial patterns displayed.

**Movie S1. Modest pattern formation by synthetic LALI circuit linked to Fig. 2C**.

Time lapse movies of coculture of Cell A and Cell B with 0 or 1000 ng/ml Dox for 72hrs. The images were taken every 2hr by Nikon AX-R confocal microscope. All time-point images were processed to show ON, OFF and NO cell regions.

**Movie S2. Synthetic pattern formation by synthetic LALI circuit coupled with cell adhesion induction linked to Fig. 3C**.

Time lapse movies of coculture of Cell A-iPcad and Cell B-iPcad with 0 or 1000 ng/ml Dox for 72hrs. The images were taken every 2hr by Nikon AX-R confocal microscope. All time-point images were processed to show ON, OFF and NO cell regions.

